# Vessel normalization and maturation promotes nanoparticle delivery to solid tumors while minimizing metastases

**DOI:** 10.1101/2023.04.27.538559

**Authors:** Mukaddes Izci, Christy Maksoudian, Filipa Gonçalves, Tianjiao Chu, Carla Rios Luci, Eduardo Bolea-Fernandez, Frank Vanhaecke, Bella B. Manshian, Stefaan J. Soenen

## Abstract

Nanoparticle delivery to solid tumors is known to be an inefficient process and various studies have tried to increase efficacy, but mechanistic and comparative studies remain scarce. Here, we use pharmacological agents to study the effect of vessel normalization or vessel disintegration on nanoparticle delivery to solid tumors. Using a multiparametric approach, we find that vessel disintegration fails to improve nanoparticle delivery and instead seems to have a limiting effect. Vessel normalization, however, improves delivery efficacy for nanoparticles ranging from 20 to 60 nm diameter. The normalization of the tumor blood vessels results in reduced hypoxia, reduced necrosis and an increase in *Plvap*^+^ *CD276*^+^ endothelial cells, which have been linked with nanoparticle delivery. Interestingly, where vessel disintegration stimulated cancer cell intravasation and associated metastases, vessel normalization impeded these processes. Together, these data reveal that, vessel normalization may be a safer and more suited approach for improving nanoparticle delivery to solid tumors, but its efficacy is limited by nanoparticle diameter and tumor parameters.

**Graphical abstract:** 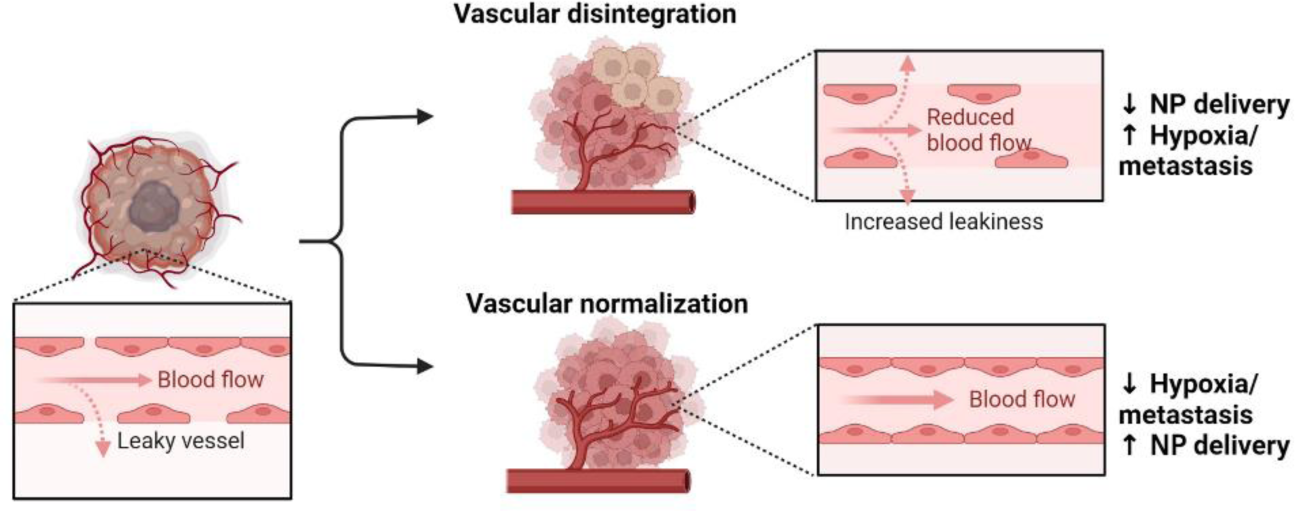

## 1 Introduction

One of the research fields that is gaining high levels of attention, is nanomedicine, being the application of nanotechnology for medical purposes. This relatively novel field is rapidly gaining much interest,(1) owing its success to the highly multidisciplinary nature of the field in itself, combining physics and chemistry expertise in nanomaterial (NM) synthesis and characterization with expertise in biology and medicine in view of functional applications. For cancer therapy and diagnosis, various NMs are already approved for clinical use, and many more are undergoing clinical trials.(2) The main application hereof lies in the use of NMs as carriers for more common chemotherapeutic agents.(1, 2) Owing to their physicochemical properties, NMs can enhance specific delivery of any pharmaceutical agent to the tumor, either passively or by stimulated (externally triggered) release.(2) Delivery of NMs to the tumor site is normally achieved in a process called enhanced permeability and retention (EPR) effect, which can occur through passive accumulation.(3) NMs are then optimized for long term circulation (*e.g*. by addition of poly-(ethylene glycol)) and their ability to extravasate into the leaky endothelium of the tumor.(4) This typically results in low levels of NM accumulation in the tumor site, thus most studies make use of active targeting ligands (*e.g*. antibodies, peptides or membranes from host cells) to increase tumor targeting.(5)

A recent meta-analysis showed that in preclinical models only 0.7% of the intravenously administered dose of NMs accumulates in solid tumours irrespective of whether this occurred via passive or active targeting.(6) Therapeutic efficacy is however not solely determined by effective NM targeting, but of course also on the efficiency of drug release. While 0.7% may not sound very convincing this value in itself is higher than the values obtained for many conventional drugs not associated with a nano-formulation.(7) Additionally, nano-formulations have been shown to dramatically enhance the time frame of tumor exposure by significantly reducing the clearance rate of the agent.(7) Nano-formulations have therefore been shown to have great clinical potential, but also have a large window of opportunity for further improvement by boosting the delivery efficacies.(8)

Various efforts have recently been undertaken to try and boost the delivery of NMs to solid tumors. One trend that has emerged is the need for personalized medicine, where depending on the physiology of the tumor of a particular individuum, this tumor would be more or less susceptible for NM therapy.(3) Where most studies to date do not classify different tumors based on their vascular properties (total blood flow and tumor vessel permeability), they frequently will observe a broad distribution in the tumor targeting ability of any administered NM. Recent studies have shown that by using an imaging contrast agent along with the therapeutic NM, a better idea of the vascular properties of every particular individuum can be obtained, based on which the decision whether or not this particular individuum would be susceptible to NM-based therapy can be made.(9)

In view of increasing NP delivery to tumours, various studies have focused on the importance of EPR and looked into ways to modify vessel permeability as a means to improve NP extravasation.(10, 11) While various reports exist that demonstrate the effect of vessel permeabilisation on NP delivery, the multifaceted nature of the tumor microenvironment (TME) and the complex interactions that are involved between the different cell types that make up the TME render it quite difficult to affect only a single parameter (vessel permeability) without affecting other aspects that would influence NP delivery efficacy. While less frequently studied, vessel normalization has also been looked into and was found to affect NP delivery efficacy.(12) As it is unclear which of both strategies would be beneficial in terms of NP delivery efficacy, the present study aims to address this question by means of a multiparametric analysis method, which was recently optimized in-house.(13) To this end, differently sized gold NPs were administered to tumour-bearing mice under conditions which either induces vessel permeabilization or vessel normalization.

## 2 Results

### 2.1 Nanomaterial characterization

The present study employs the use of gold nanoparticles (Au NPs) as model systems to study NP distribution. Au NPs were selected owing to the high robustness and reproducible synthesis of NPs with tight control over size and surface properties. The use of Au furthermore enables sensitive quantification of tumor-delivered NPs by means of inductively coupled plasma mass spectrometry (ICP-MS), while darkfield-based imaging together with image-based cytometry can be used to evaluate cellular NP uptake at the single cell level and make distinctions between the cell types in the TME that play a role in NP uptake. Due to these aspects, Au NPs have also been widely used in NP biodistribution studies,(6, 14) resulting in a large literature portfolio that can be used for interpreting the data obtained. The NPs used have a theoretical size of 10, 20, 40, 60 and 80 nm in diameter (defined as Au_10_, Au_20_, Au_40_, Au_60_ and Au_80_, respectively) and are coated with poly(metacrylic acid) and 2 kDa methoxy-poly(ethylene glycol) (PEG). These NPs are new batches of the same ones as used in our previous work, where we looked at the effect of NP size on tumor delivery efficacy, and can be seen as reference values.(13) NP core sizes were evaluated using transmission electron microscopy (TEM), the hydrodynamic diameter and surface charge were measured using dynamic light scattering and zeta-potential measurements while colloidal stability of the NPs in serum-containing media was evaluated using nanoparticle tracking analysis.

**Figure 1** provides an overview of NP characteristics, revealing a narrow size distribution, good colloidal stability and a diameter that matches the theoretical diameter (provided by company that synthesized them). All NPs possessed a negative surface charge and were on average 20 nm larger in size due to the presence of the PEG chains and immobile unit of the solvent ions. Interestingly, comparing the values to the ones in our previous study revealed a high interbatch reproducibility as NP properties are highly similar. To study the effect of tumor blood vessel modifications, the experimental setup was performed as schematically illustrated in **Figure 2**.

**Figure 1.**
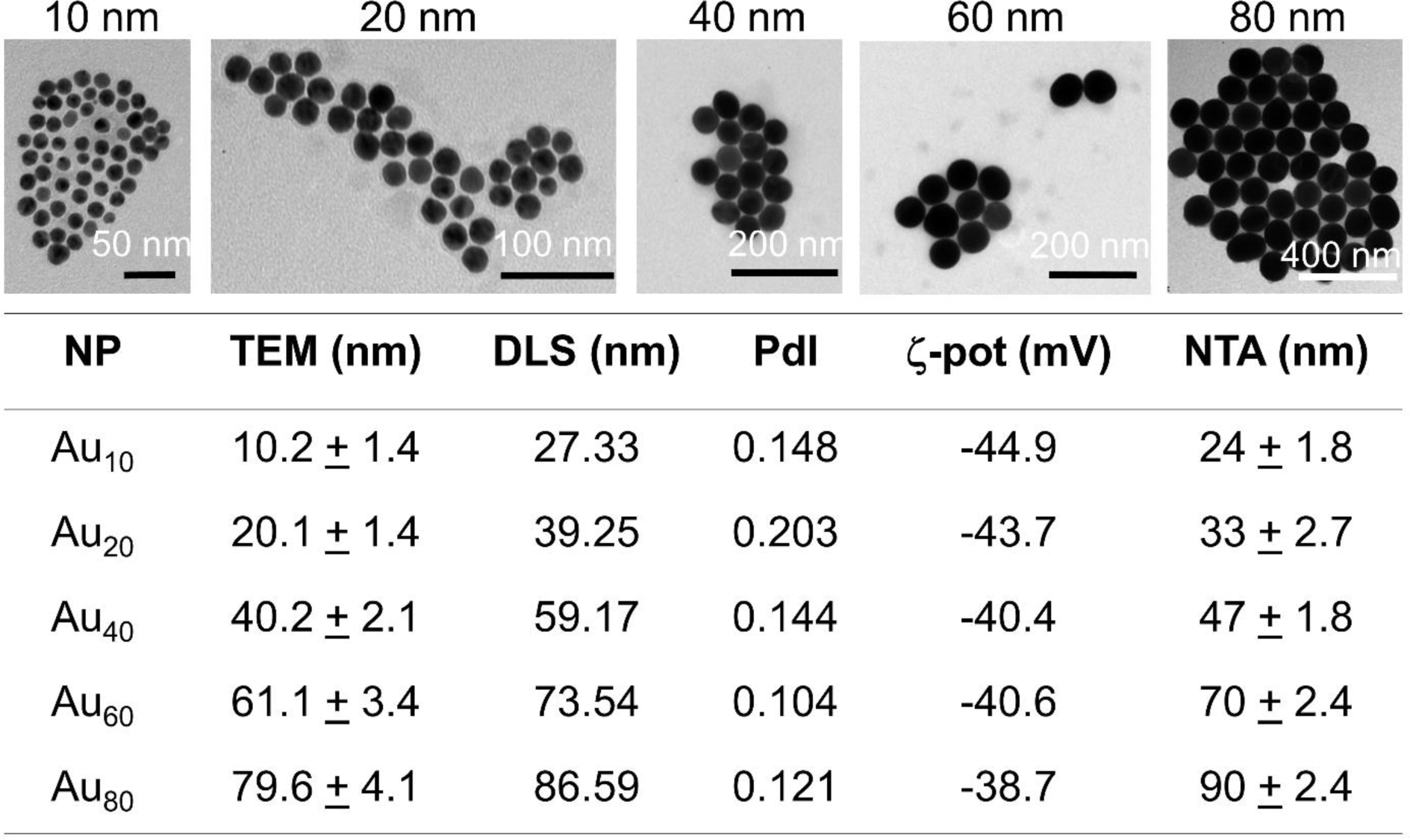
Nanoparticle characterization data. Representative transmission electron micrographs are given for the 5 differently sized gold nanoparticles that are used in this study. The table below show various parameters for every nanoparticle used. The first parameter is the core diameter (determined by TEM) and analysed by measuring 100 NPs over different images. The second parameter is the hydrodynamic diameter in aqueous environment (determined by DLS in PBS). The third parameter is the polydispersity index (PdI), determined simultaneously by DLS which indicates colloidal stability of the NPs in PBS. The fourth parameter is the ζ-potential, which is the NP surface charge, as measured in PBS. The fifth parameter is the colloidal stability of the NPs in high levels of serum (as determined by NTA in 50% FBS-containing PBS).

**Figure 2.**
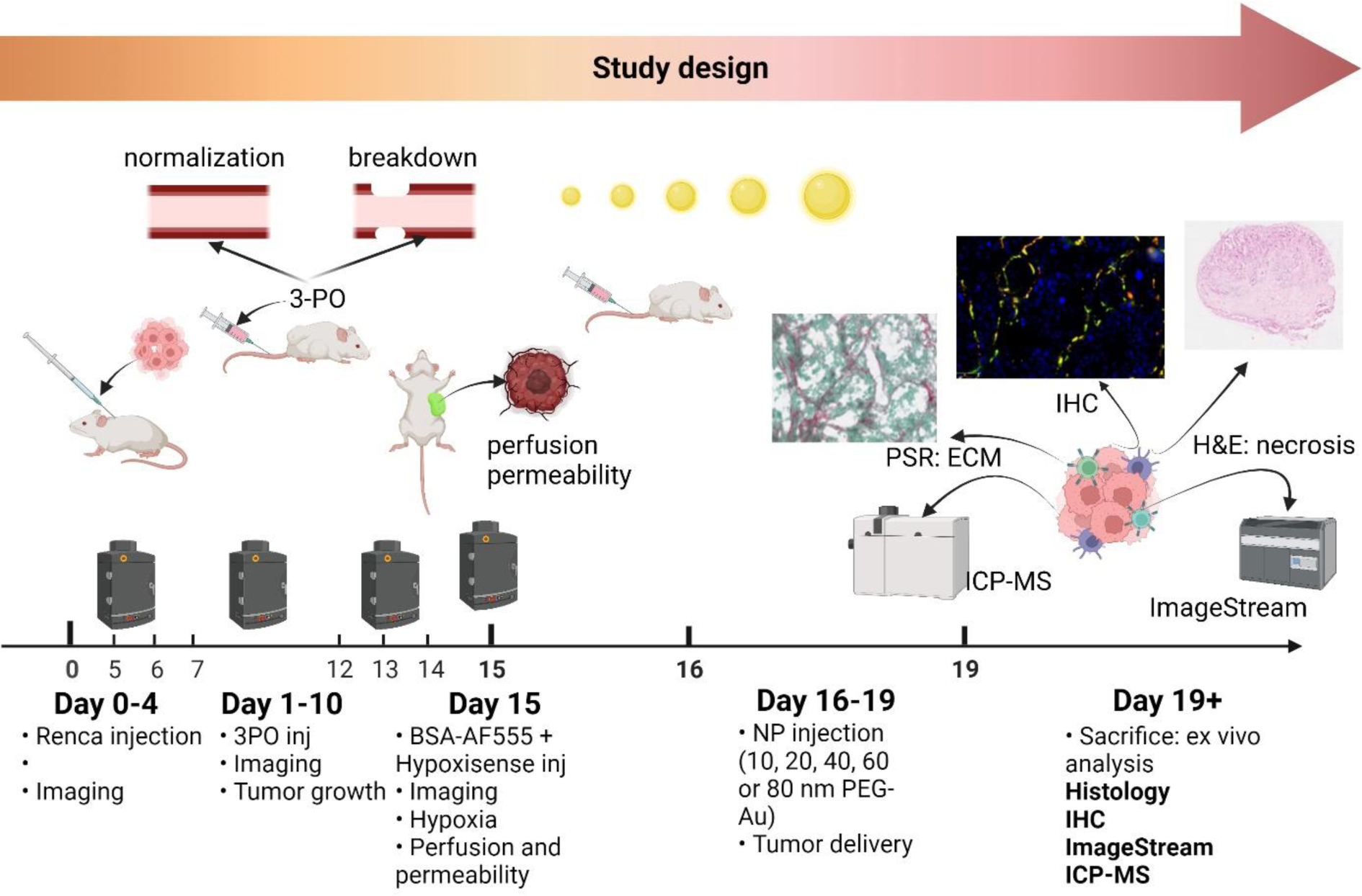
Schematic overview of the study design. Upon subcutaneous administration of luminescently-tagged Renca cells, non-invasive imaging was performed at days 4, 7, 10 and 14 to monitor tumor growth. At day 5,6,7 following Renca cell injection, tumor-bearing animals received a bolus of 3-PO every other day at either low (25 mg/kg) or high (75 mg/kg) dose. This was repeated at days 12, 13 and 14. At day 15, AF555-tagged BSA and Hypoxisense 680 were administered intravenously and optical imaging was performed to determine relative perfusion and permeability of tumor-associated blood vessels as well as measure differences in tumor-related hypoxia levels. At day 15, animals received 100 µL of PBS with Au NPs (150 µg Au/mouse, either 10, 20, 40, 60 or 80nm diameter in size) by intravenous administration. The NPs were then left to circulate for 72 hrs, after which time we had observed a complete absence of the NPs in the blood (see manuscript Izci *et al*. for more details),(13) after which the animal and the tumors were analyzed *ex vivo* for potential toxicity, NP tumor delivery efficacy, and tumor-associated parameters as indicated in the image.

### 2.2 Practical setup of the experimental design

Firefly luciferase-expressing renal carcinoma cells (Renca) were used in this study, as a well-developed and commonly used syngeneic tumor model. The Renca cells were chosen as they had been previously used by our group for biodistribution studies and therefore allowed us to compare the data obtained with those from earlier work. It is important to note that we have opted for a syngeneic model system as for our investigation, we wanted a tumor with complete tumor microenvironment (TME) as the immune cell components of the TME play a major role in NP uptake. Any human cells would therefore be quite limiting as these would require immunodeficient mice which would lack major components of the TME. On the other hand, it is important to note that the murine TME is not an exact copy of the human TME and this study aims to shed insight into the precise components affecting NP tumor delivery, which in turn can be exploited in clinical settings. Another important factor is that the Renca cells were used as a subcutaneous model system, and were not implanted orthotopically. This inherently means that the TME will be affected, which in turn will influence NP delivery efficacy. The subcutaneous model was chosen however as for preclinical research it is the most commonly used model system, and in our hands, the readouts of hypoxia, tumor perfusion and tumor permeability using optical imaging enables high-throughput analysis but is limited to subcutaneous models due to problems in fluorescence intensity readings at larger depth. Furthermore, as shown in this and other studies, while the same tumor cells were used in the same animal model, the TME composition varies greatly and no two tumors can be called identical, despite being the same ‘model’. Therefore, this study sets out to carefully characterize the TME and the role of different components in tumor targeting which will lead to more generic insights on how to improve tumor targeting.

One week following tumor grafting, the animals are treated with 3-(3-pyridinyl)-1-(4-pyridinyl)-2-propen-1-one) (3PO) every other day for a total period of 10 days. The high glycolytic activity of tumor cells and tumor-associated endothelial cells has made inhibition of glycolysis by blocking the glycolytic activator 6-phosphofructo-2-kinase/fructose-2,6-biphosphatase 3 (PFKFB3) a powerful treatment option.(15) 3PO is an effective PFKFB3 blocker 3-(3-pyridinyl)-1-(4-pyridinyl)-2-propen-1-one) and its use has been studied in greater detail, finding, that depending on the dosage used, this could either result in tumor vessel normalization (25 mg/kg; 3PO_lo_) or disintegration (at higher dose: 75 mg/kg; 3PO_hi_).(16)

PFKFB3 activates glycolysis by synthesizing fructose-2,6-bisphosphate (F2,6P2), an allosteric activator of phosphofructokinase-1 (PFK-1), a rate-limiting enzyme of glycolysis.(16) Literature reports have shown that a high maximum tolerable dose of the PFKFB3 blocker 3PO inhibits tumor growth,(17) and a phase I clinical trial on advanced solid malignancies is being carried out.(18) One important consideration is that apart from cancer cells, tumor endothelial cells (TECs) are also highly glycolytic.(15) Tumor vessels are structurally and functionally highly abnormal,(19) rendering them hypoperfused, depriving cancer cells from oxygen and nutrients, thereby creating a hostile environment from where cancer cells attempt to escape.(20) In turn, the predominance of neoangiogenesis results in the rapid generation of leaky blood vessels rather than blood vessel maturation, promoting cancer cell dissemination. Anti-angiogenic strategies that further aggravate these tumor vessel abnormalities bear the risk of increasing metastasis, while impairing chemotherapy.(20) In contrast, tumor vessel normalization offers opportunities to reduce metastasis, while improving chemotherapy by reducing interstitial pressure.(19, 21, 22) The effect of tumor vessel normalization or disintegration on NM delivery has only been scarcely described. As our previous results indicated that permeability of the blood vessels did not appear to be positively correlated with improved NP delivery,(13) and various literature sources indicate increased NP delivery efficacy with either vessel normalization or increased permeabilization,(10, 12, 23) we set out to investigate the influence of tumor vessel modification on NP delivery and whether one of either methods would be more beneficial.

### 2.3 Efficacy of tumor vessel modulation

As a first test, the effect of 3PO treatment on tumor growth and necrosis was assessed (**Figure 3a,b**). While 3PO_lo_ displayed no significant effects on tumor growth compared to untreated control animals, 3PO_hi_ resulted in significantly reduced tumor growth, which is in line with literature reports.(16, 17, 24) These effects likely stem from a combination of two factors, one being the direct inhibition of PFKFB3 in the highly glycolytic cancer cells and therefore impeding cancer cell proliferation.(16) Alternatively, highly glycolytic TECs can also be affected as TECs have been found to increase glycolysis to meet their high demands of biomass and ATP synthesis for rapid proliferation and migration.(16) Specifically targeting TEC glycolysis has been found to inhibit angiogenesis and at higher levels (similar to the concentrations used in this study), 3PO has been shown to cause tumor vessel disintegration.(16) On the level of necrosis, 3PO_lo_ and 3PO_hi_ have contrasting effect, where they are resulting in a significant decrease and a significant increase, respectively. These data hint towards an improved delivery of nutrients and oxygen to the tumor inflicted by 3PO_lo_ treatment. Using high-throughput optical imaging, the perfusion and permeability of the tumor was evaluated. To this end, fluorescently labeled bovine serum albumin (AF555-BSA) was administered intravenously to the mice and the kinetics of BSA distribution were evaluated for 90 minutes (**Figure 3c-e**). Perfusion and blood vessel permeability were then determined as relative units compared to the skull base, where only minimal leaky blood vessels would be expected. In terms of perfusion, measured as the total BSA level at the tumor level 5 minutes after administration, revealed no significant effect of 3PO_lo_, while 3PO_hi_ caused a significant reduction. The latter is in line with the increased necrosis observed as the poorly perfused tumor would result in poor nutrient availability. This was further confirmed by looking into the level of hypoxia (**Figure 3f-h**). 3PO_lo_ significantly reduced hypoxia levels, while this was minimally elevated in 3PO_hi_-treated animals. These data further indicate the improved delivery of oxygen to the tumor by 3PO_lo_, while 3PO_hi_ did not seem to have a major role in oxygen availability. The latter may be attributed to intrinsic properties of the tumor model used, where Renca tumors are known to be relatively fast-growing and aggressive tumors, that suffer from high levels of hypoxia and large necrotic cores.(25)

**Figure 3.**
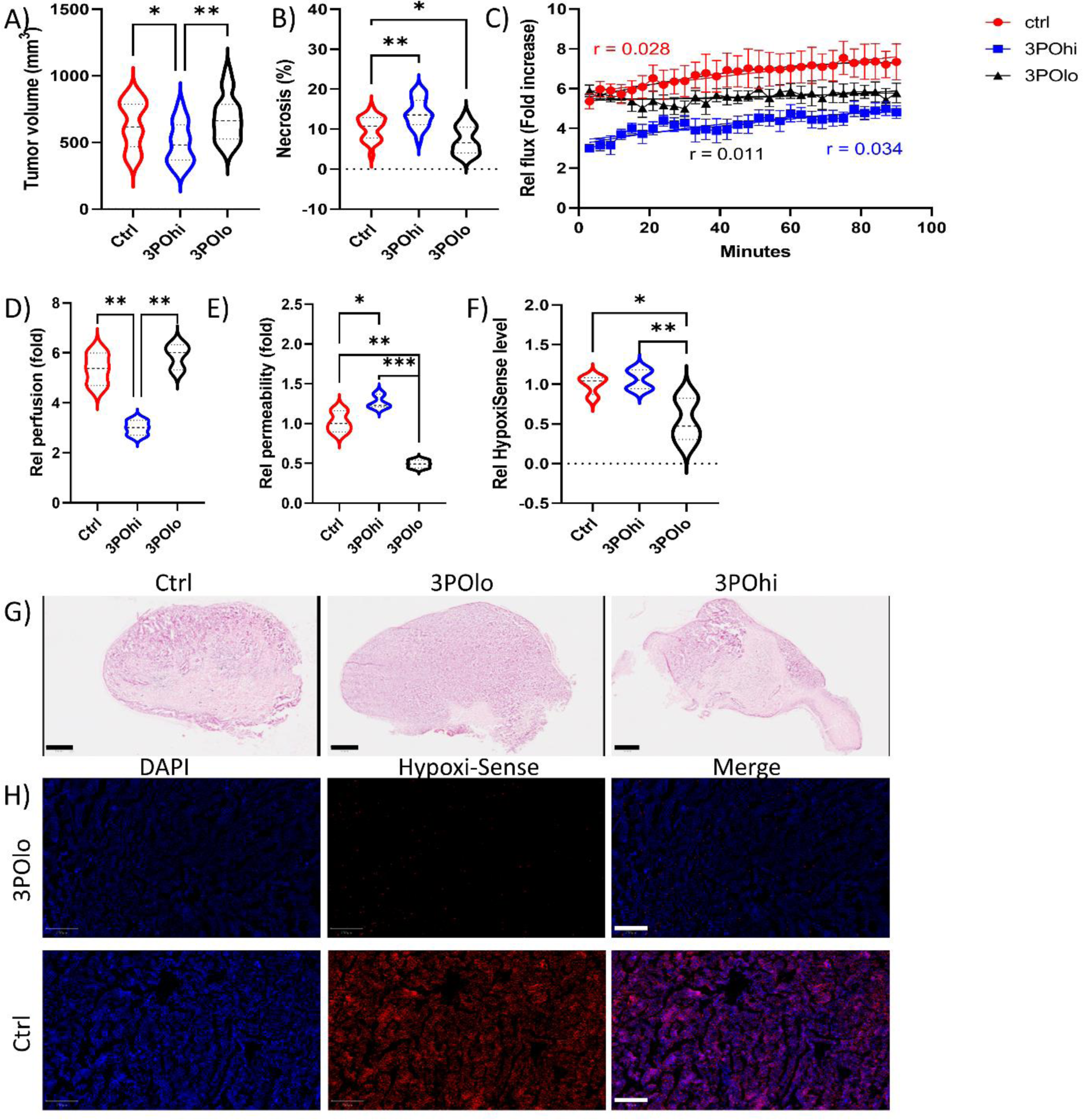
The effects of 3PO treatment on tumor physiology. a,b) Violin plots of the respective indicated parameter for every group of animals (*n* = 10). a) The tumor volume, expressed in mm³ and measured by caliper measurements immediately prior to tumor resection. b) The area of necrotic tissue in the tumor as determined by H&E staining relative to the area of the entire tissue slice. c) Histogram displaying the relative fluorescence signal of AF555-BSA as a function of time in the 3 conditions indicated (control, 3PO_hi_ (75 mg/ml); 3PO_lo_ (25 mg/ml)). Data are expressed as the fold difference of the tumor-associated signal over the signal derived from the base of the skull of the same animal as an internal reference. Data are presented as mean ± SEM (*n* = 10 per condition). d-f) Violin plots of the respective indicated parameter for every group of animals (*n* = 10). d) Perfusion in the tumor, expressed as the fold difference in AF555-BSA signal in the tumor compared to the skull 5 minutes following contrast agent administration. e) Permeability of the tumor blood vessels, expressed as the fold difference in AF555-BSA signal in the tumor compared to the skull from 5 until 90 minutes long, where the slope of the curve indicates permeability. f) Hypoxia levels expressed as the signal intensity of HypoxiSense 680-treated animals. The data are presented as relative values compared to control tumor bearing animals without 3PO treatment (average control value = 1). g) Representative H&E-stained tumor sections obtained from differently treated animals, demonstrating the influence of 3PO on tumor necrosis. Scale bar: 1 mm. h) Representative fluorescence images of tumor sections obtained from control animals and 3PO_lo_ animals treated with HypoxiSense 680 (red) and stained with DAPI nuclear stain (blue). Scale bar: 50 µm. Significant differences in all violin plots are indicated where appropriate (n = 10; *: p < 0.05; **: p < 0.01; ***: p < 0.001).

This was further evaluated by looking into different tumor-associated parameters of all animals used in this study for NP delivery and treated with 3PO or not (**Figure 4**).

**Figure 4.**
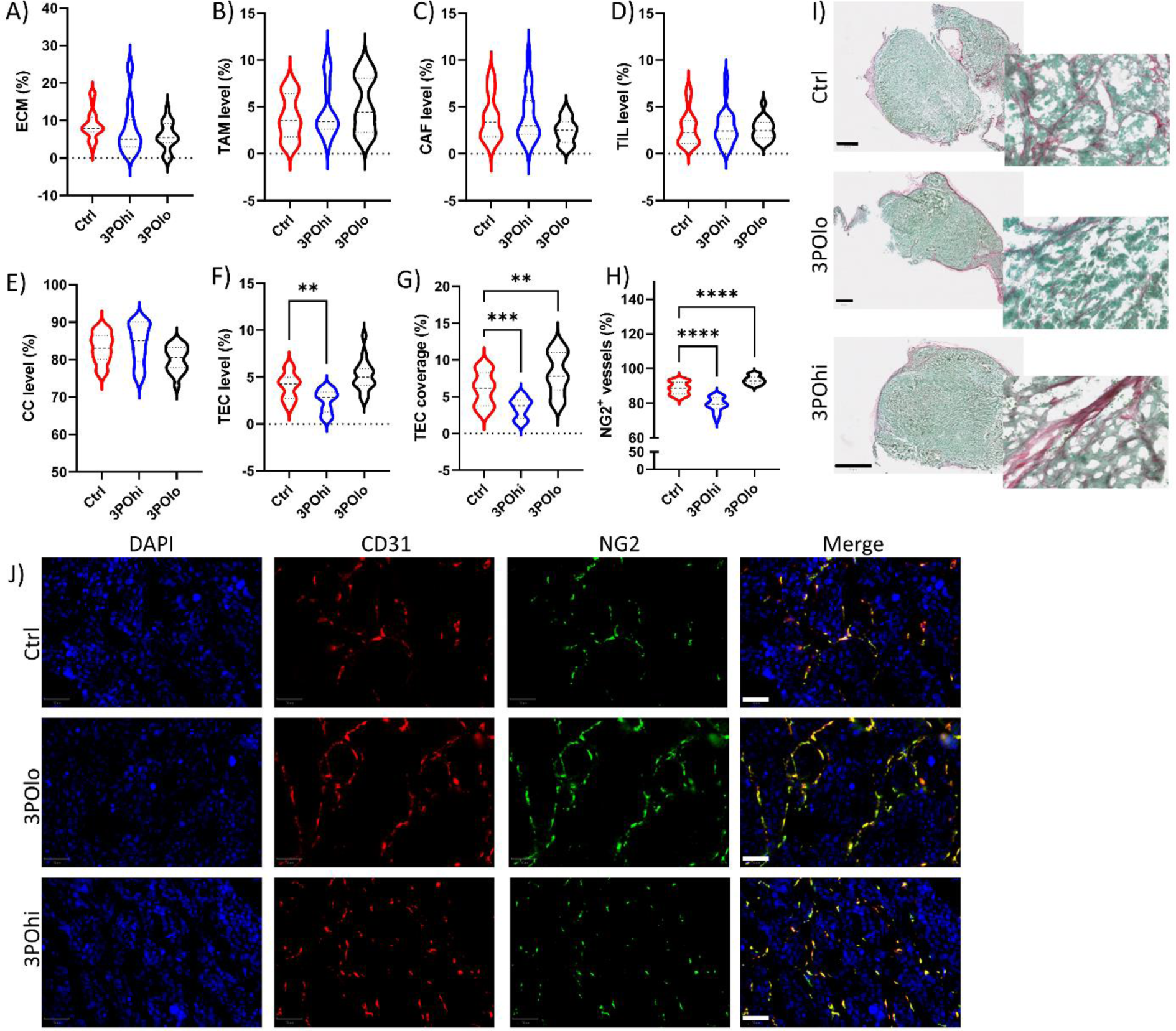
The effects of 3PO treatment on TME composition. A-h) Violin plots of the respective indicated parameter for every group of animals (*n* = 25). a) The area positive for ECM in the tumor as determined by Picrosirius Red staining relative to the area of the entire tissue slice. b**-f)** Relative level of b) TAMs, c) CAFs, d) TECs, e) TILs and f) cancer cells (CC) expressed relative to the total number of cells as determined by ImageStreamX Mark II analysis. g,h) The relative level (%) of g) tumor area occupied by CD31+ endothelial cells, h) CD31+ blood vessels covered by NG2. I,j) Representative images of i) PSR and FastGreen-stained tumor sections and j) immunofluorescence staining of tumor tissues against endothelial cell marker CD31 (red) and pericyte marker NG2 (green). Scale bars: 50 µm. Significant differences in all violin plots are indicated where appropriate (*n* = 25; *: p < 0.05; **: p < 0.01; ***: p < 0.001).

The tumors themselves were analyzed for a variety of additional parameters including the density and extent of the extracellular matrix, the level of tumor-associated macrophages (TAMs), cancer-associated fibroblasts (CAFs), tumor-infiltrating lymphocytes (TILs), TECs, cancer cells (CC), as well as the relative tumor area occupied by blood vessel and tumor blood vessel maturity expressed as their coverage by pericytes (**Figure 4**). Data across all groups revealed that while individual values for particular parameters could vary, most parameters did not seem to be affected by 3POlo or 3POhi treatment. These data suggest that for the NP delivery studies, individual tumors had variable levels for all of these parameters, but the average of the group of animals did not significantly differ for most parameters. However, in line with expectations and the data displayed in Figure 3 indicating changes in perfusion and tumor vessel permeability, the total number of TECs, as well as the area of the tumor covered by blood vessels as well as blood vessel maturity were significantly affected by 3PO_lo_ and 3PO_hi_ treatments. In both cases, 3PO_hi_ treatment reduced TEC coverage and blood vessel maturity, while 3PO_lo_ treatment increases both parameters (**Figure 4g,h**).

Taken together, these data indicate the 3PO_hi_ clearly causes tumor vessel disintegration, which results in reduced tumor growth, but also causes elevated hypoxia and necrosis caused by lower tumor vessel perfusion levels. For 3PO_lo_, the situation appears to be a bit more complex as most parameters suggest that 3PO_lo_ resulted in tumor vessel normalization, but this did not lead to significant changes in view of tumor vessel perfusion levels. Overall, the outcome of antiangiogenic strategies relies on several conditions to be met to determine whether the treatment elicits vessel normalization of vessel pruning.(26) Firstly, the vascular density level must be sufficiently high for the treatment to work. As excessive pruning would result in hypoxia and here the opposite result is obtained, we can assume that no excessive pruning is induced, but rather that vessel normalization is promoted. Secondly, there has to be a sufficiently high level of non-compressed blood vessels. As ECM density is typically very high in solid tumors and is also correlated with tumor growth,(27) the high pressure caused by the thick ECM can collapse tumor blood vessels. In our model, untreated mice already display high perfusion levels, which can be significantly reduced upon tumor vessel disintegration. A third condition relies on the need for pericyte-covered blood vessels, thereby enabling further maturation and fortification of blood vessels. The *ex vivo* analysis of tumor blood vessel shows clear pericyte-covered TECs, which is further increased upon 3POlo treatment. As all conditions are met, the apparent lack in tumor vessel perfusion likely stems from the relatively good tumor vessel growths in the Renca tumor model that was used in this study. While renal adenocarcinomas typically respond well to antiangiogenic therapies due to mutation-induced vascular endothelial growth factor (VEGF) upregulation(28) as well as stabilization of the alpha subunit of hypoxia-inducible factor (HIF), resulting in chronic hypoxia, Renca cells themselves do not express the latter mutations.(29) Taken together, 3PO_lo_ does appear to cause slightly, but not significantly elevated increases in tumor perfusion, but most importantly, results in significantly higher levels of vessel maturation, reduced hypoxia and reduced vessel permeability.

### 2.4 Evaluation of nanoparticle delivery to solid tumors

To unravel the effect of 3PO_lo_ or 3PO_hi_ treatment on NP tumor delivery efficacy, tumor bearing mice treated with 3PO_hi_, 3PO_lo_ or vehicle were given a single bolus of Au NPs of either 10, 20, 40, 60 or 80nm diameter. After 72 hrs, animals were sacrificed and detailed studies were performed in accordance with our previous report.(13) Blood biochemistry and histopathological analysis of all major organs indicated no toxicity induced by either the NPs themselves or 3PO treatments (**Supplementary Figure S1**). When looking at the overall value of tumor-associated NPs, no clear size-dependent differences were observed for control, 3PO_hi_ or 3PO_lo_ treated animals (**Figure 5a-c**). The lack of any clear NP size-dependent tumor delivery efficacies is in line with our previous study, and averaged approximately 0.7% of the originally injected dose of NPs, which is in line with the meta-analysis on tumor delivery published in 2016. One clear observation from these initial data is that for 3PO_hi_ and 3PO_lo_ groups, the average NP delivery appeared to be reduced and increased, respectively, apart from Au_10_ NPs, where none of the treatments appeared to have any significant effects. To verify the influence of 3PO treatment on NP delivery, the % of NPs delivered is presented for every NP size separately (**Figure 5d-h**).

**Figure 5.**
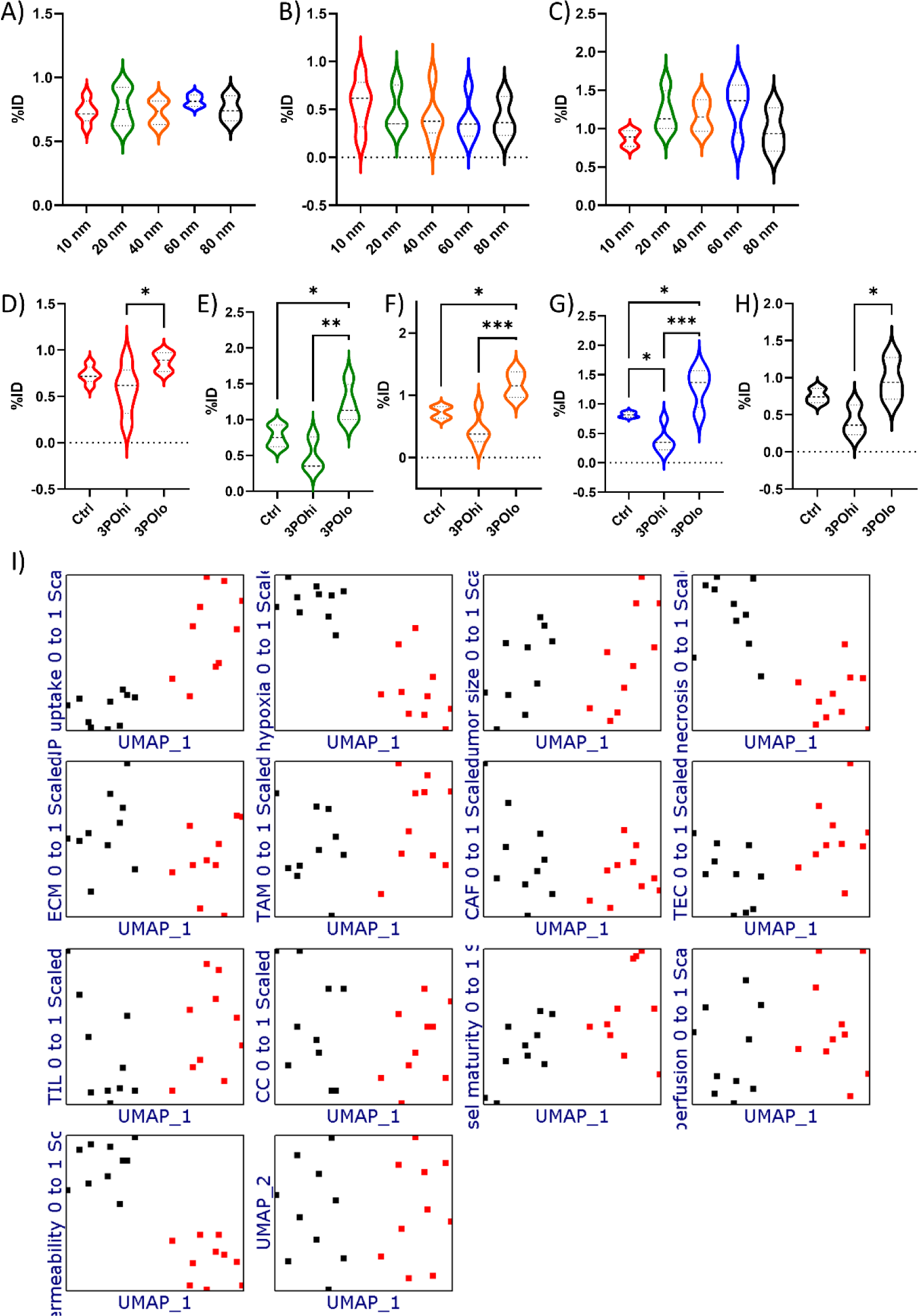
Overview of NP delivery efficacy and the role of 3PO treatment. **A-c)** Violin plots showing the NP delivery efficacy for every group of NPs expressed as the % of tumor-associated NPs relative to the originally administered amount. The data is shown for every size of NP used for the 3 main conditions, being **a**) untreated control mice, **b**) 3PO_hi_ treated mice and **c**) 3PO_lo_ treated mice. **d-h**) Violin plots showing the NP delivery efficacy for every group of NPs expressed as the % of tumor-associated NPs relative to the originally administered amount. The data is shown for control (untreated) mice, 3PO_hi_ and 3PO_lo_ treated mice for d) Au_10_ NPs, e) Au_20_ NPs, f) Au_40_ NPs, g) Au_60_ NPs and h) Au_80_ NPs. Significant differences in all violin plots are indicated where appropriate (*n* = 15; *: p < 0.05; **: p < 0.01; ***: p < 0.001). i) Representative UMAP plots for each and every tumor-associated parameter determined (as displayed in Figures 3 and 4) as a function of UMAP coordinates and displayed for Au_40_ NPs for untreated (black dots) and 3PO_lo_ (red dots) groups. For every parameter, the values were first rescaled to a linear 0-1 scale with 0 being the lowest value for that parameter across all animals and 1 being the highest value for that parameter across all animals. Every single dot shown reveals a separate animal and indicates the specific value of the animal on its Y-axis, while for every animal, its position on the X-axis does not change and thus, values for all parameters can be directly compared. To determine whether a particular parameter promotes or inhibits NP delivery efficacy, the red and black groups should be separated on the Y-axis for the parameter. If the two groups can be separated from one another based on these parameters, then they will display distinct UMAP features and will also be separated on the X-axis.

The data reveal a clear significant increase in tumor delivery efficacy for Au_20_, Au_40_ and Au_60_ upon 3PO_lo_ treatment, while 3PO_hi_ led to significantly reduced delivery of Au_60_. The latter may seem somewhat surprising as EPR is founded on the principle that NPs can extravasate into the tumors owing to gaps between endothelial cells lining the blood vessels. Many studies have also reported on how exploiting or promoting the leakiness of tumor blood vessels can promote NP delivery.(10, 30, 31) However, the concept of EPR has become more of a debate, which is particularly driven by the apparent lack of clinical translation of EPR.(32) Recent studies have demonstrated that the EPR effect results in less than a 2-fold increase in tumor-restricted NP delivery compared with delivery to normal major organs.(33) NP delivery through the EPR effect is influenced by a variety of factors, including blood flow, interstitial fluid pressure, density of the extracellular matrix and the presence of occluded or embolized tumor blood vessels.(11) Previous studies have shown that most tumors possess high levels of interstitial fluid pressure, and that the level of this correlates with increased vessel permeability and hyperperfusion.(34) For 3PO_hi_, both these conditions were observed, which would explain the reduced delivery for most NPs, reaching significant levels for Au_60_. One exception in our study are the Au_10_ NPs, which do not appear to be much affected by either 3PO_hi_ or 3PO_lo_ treatment. Smaller NPs (< 10 nm) have been reported to effectively permeate inside tumors via multiple mechanisms, including trans- and extravascular transport.(11) Given the minor effect of 3PO_lo_ on tumor perfusion, this may explain why the smallest NPs were only minimally affected.

For 3PO_lo_-treated animals, the average level of tumor-delivered NPs is higher than for 3PO_hi_- or PBS-treated animals. For NPs sized between 20 and 60 nm, this is even a significantly elevated level of NPs (**Figure 5e-g**). For the largest NPs, the lower efficacy in tumor delivery enhancement may seem somewhat logical, as their larger size will limit the transfer through the less frequently occurring tumor vessel endothelial gaps. However, the smaller sized NPs would be considered to be well suited for improved therapy, as the key initial study by the group of Rakesh Jain confirmed vessel normalization resulting in increased delivery of 12nm diameter gold NPs while they saw no effect in bigger sized NPs.(35) Further studies revealed that the same strategy for tumor vessel normalization, did also result in increased tumor delivery of “medium-sized” NPs, being 20 and 40nm diameter.(36) The apparent differences between various studies on vessel normalization are unfortunately difficult to relate to a single parameter. Vessel normalization is a complex multifactorial process which in itself may also influence other components of the TME, such as the density of the ECM or the relative level of tumor-associated myeloid cells. Furthermore, the tumor type used in the study itself, its location for grafting and the time window in which the NPs will be given will all influence perfusion and vessel permeabilisation parameters. In our study, vessel normalization seems to only exert a minimal impact on tumor perfusion, which may stem from the fact that the Renca model that was chosen is already well perfused.

### 2.5 The influence of vessel normalization on NP delivery efficacy

When looking at the individual parameters and their distinction for PBS- and 3PO_lo_-treated animals, statistical modelling was performed, where all parameters were first modeled on a scale from 0 (lowest value) to 1 (maximum value) in order to avoid skewing the data towards parameters with higher numerical values. Then Uniform Manifold Approximation and Projection (UMAP) dimensionality reduction analyses were performed that enables multiparametric analysis to be grouped and expressed over two dimensions to reveal similarities or distinctions between the two groups. For ease, the PBS-treated group was indicated in black, while the 3PO_lo_ group was indicated in red. The data shown in **Figure 5i** is a representative plot for Au_40_ NPs, which revealed that the PBS- and 3PO_lo_ group could be easily separated from each other is evidenced on the X-axis spreading. When looking at the individual parameters (Y-axis spreading), the parameters that show most distinct differences are increased NP delivery to the tumor, decreased hypoxia, decreased necrosis, and decreased vessel permeability upon 3PO_lo_ treatment. The level of hypoxia, necrosis and vessel permeability therefore seem to play the key roles in determining the efficacy of 3PO_lo_ treatment on NP delivery.

In an editorial by Andre Nel and colleagues on NP delivery, the question was asked whether NP extravasation is primarily a process of NPs slipping through abnormal fenestrations or gaps between adjacent endothelial cells or is the major access route transendothelial transport pathways that assist tumor growth and nutrition?(37) Recent work by the Warren Chan group utilized advanced murine model systems, the so-called “zombie mice” in which active transport mechanisms across the endothelial barrier were halted and they observed that passive transfer of NPs delivered through endothelial gaps played only a minimal role in the amount of NPs delivered to the tumor.(14) Further work by the same group found that NP uptake mainly occurred through active transendothelial transport, which involved the use of specialized endothelial cells, dubbed N-TEC cells.(38) In our previous study, we were able to confirm their finding, and observed that tumors with higher levels of N-TECs gave rise to increased levels of tumor-located NPs.(13) Here, we set out to see whether 3PO_lo_ gave rise to higher levels of N-TECs which hereby could result in higher levels of NP delivery efficacy. To study this, *Plvap* and *CD276* expression levels were evaluated using RNAScope technology on 5 control tumors and 5 tumors from 3PO_hi_ or 3PO_lo_-treated animals, that resulted in significantly different NP delivery efficacy, despite most physiological parameters apart from hypoxia, necrosis and vessel permeability being unchanged (**Figure 6a-e**). The markers *CD276* and *Plvap* were selected based on the study by Kingston and colleagues, who described these two genes as being significantly upregulated in N-TECs compared to other endothelial cells,(38) and which were successfully used in our previous study.(13) The data obtained by the RNAScope analyses support our hypothesis and revealed that tumor vessel normalization results in higher levels of N-TECs, which is linked to a clear correlation with NP delivery. As vessel normalization mediates NP delivery through increased N-TECs levels, and NP size-dependent differences are observed, we postulated that this is due to a favorable size of the NPs for N-TEC interactions as variations in 3PO_lo_ efficacy seems unlikely. This matches previous reports indicating that cellular interactions are size-dependent and have an optimal size of around 40-50 nm.(39, 40)

**Figure 6.**
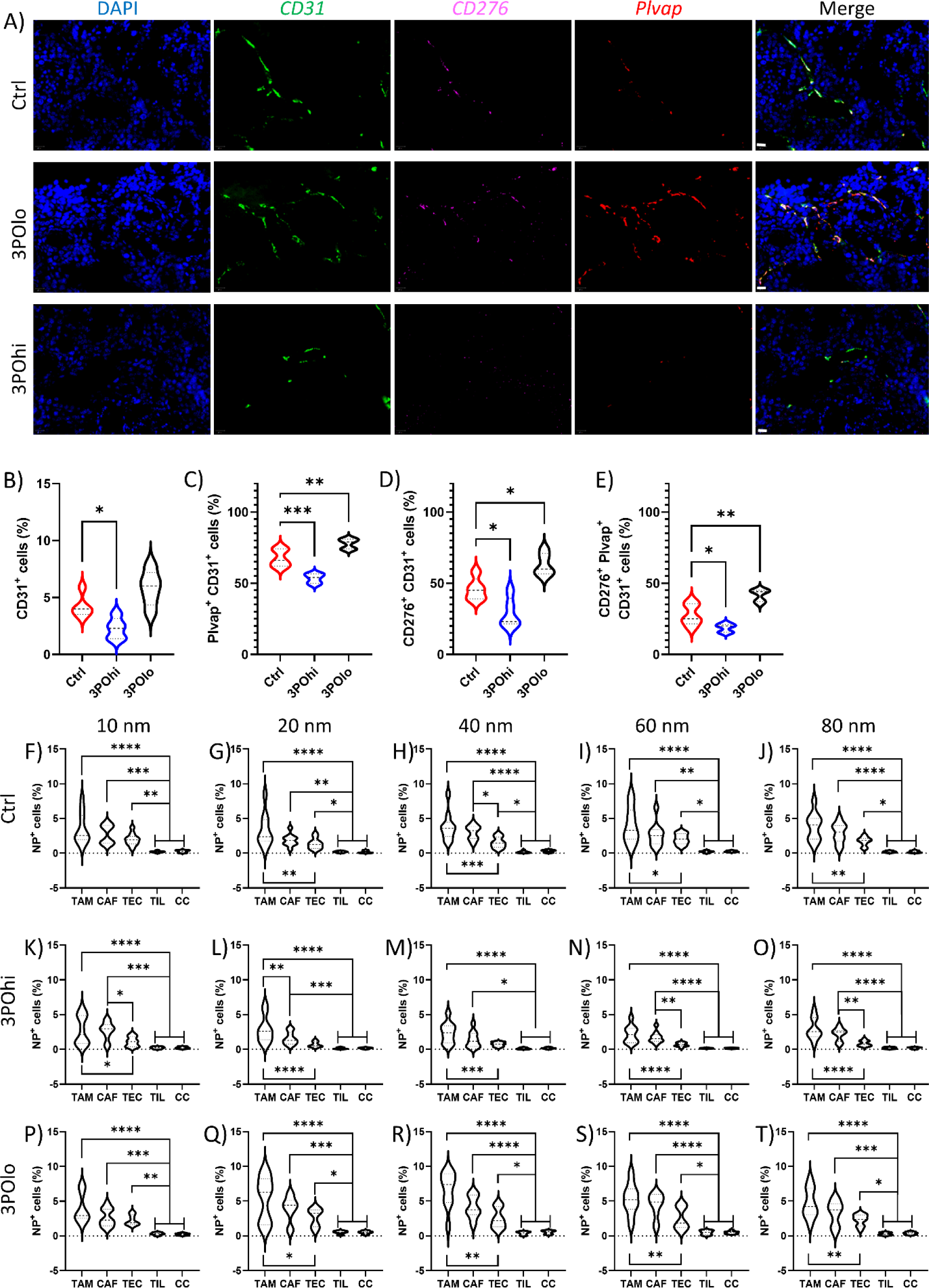
The effect of 3PO treatment on NP delivery to solid tumors and cancer cells. **a)** Representative RNAscope images of tumor sections stained against CD31, CD276, Plvap mRNA and counterstained with DAPI nuclear stain. Scale bars = 20 μm. The top row is from a control tumor, the middle row from a tumor treated with 3PO_lo_ and the bottom row from a 3PO_hi_-treated tumor. **b-e**) Violin plots of quantified RNAscope data expressed as the % of b) CD31+ cells relative to the entire cell number, c) Plvap^+^ CD31^+^ cells relative to the total CD31^+^ cell population, d) CD276^+^ CD31^+^ cells relative to the total CD31^+^ cell population and e) Plvap^+^ CD276^+^ CD31^+^ cells relative to the total CD31^+^ cell population. Statistically significant differences between 3PO_hi_ or 3PO_lo_-treated groups and controls are indicated where appropriate (*n* = 5; *: p < 0.05; **: p < 0.01; ***: p < 0.001). **f-t**) Violin plots showing the relative percentage of NP^+^ TME-related cell types expressed as the % of NP^+^ cells as determined by ImageStreamX Mark II expressed relative to the total gated cell type. The data are shown for control tumors (top row, f-j), 3PO_hi_-treated tumors (middle row, k-o) and 3PO_lo_ treated tumors (bottom row, p-t) for the differently sized NPs f,k,p) Au_10_ NPs, g,l,q) Au_20_ NPs, h,m,r) Au_40_ NPs, g,n,s) Au_60_ NPs, j,o,t) Au_80_ NPs. Statistically significant differences between any cell types per condition are indicated where appropriate (*n* = 5; *: p < 0.05; **: p < 0.01; ***: p < 0.001).

### 2.6 Evaluation of NP delivery to tumor cells

The increased delivery of NPs to the tumor region upon 3PO_lo_ treatment for NPs sized between 20 and 60nm diameter furthermore raises the question where in the tumor the NPs end up, and to what extent the NPs can reach actual tumor cells and not the TME. Therefore, we analyzed tumor-specific delivery of NPs at the single cell level using an image-based cytometry approach which was described in our previous work.(13) In short, tumors were rendered into single cell suspensions and these were stained with a mixture of antibodies to identify TME subpopulations, being cancer cells (CD45^-^CD24^+^), tumor-associated macrophages (TAMs; F4/80^+^), cancer-associated fibroblasts (CAFs; CD45^-^CD90.2^+^), tumor-associated endothelial cells (TECs; CD144^+^) and tumor infiltrating lymphocytes (TILs; CD45^+^CD3^+^). The data reveal similar trends in NP delivery to different TME subpopulations, where TAMs and CAFs overall have the highest level of NP-containing cells, followed by TECs and minimal levels for TILs and CCs (**Figure 6f-t**). These findings are perfectly in line with our previous study and literature reports, where TAMs are known to be the major cell type involved in NP uptake in the tumor environment.(41) The number of cancer cells seems very low, but this is partly driven by the TME composition of the syngeneic tumor model, where the vast majority of cells recovered are actual cancer cells. The low targeting efficacy towards cancer cells therefore ultimately reaches a considerable number of actual cells; which may even exceed TAMs, but relatively, the respective contribution is far lower.

Next, we observed clear effects of both 3PO_lo_ and 3PO_hi_ treatment on NP delivery to different subpopulations (**Figure 7 a-k, Supplementary Figure S2**). In line with the overall tumor delivery, 3PO_lo_ resulted in improved delivery of NPs while 3PO_hi_ resulted in reduced NP delivery levels. The effects were also mostly apparent for Au_20_, Au_40_ and Au_60_, while this was less extensive for Au_10_ and Au_80_. While apart from TILs, all cell subpopulations seemed to be affected by 3PO treatment, cancer cell-specific delivery was improved for Au_20_, Au_40_ and Au_60_ NPs upon treatment with 3PO_lo_. To validate whether cancer cells could be more specifically targeted upon 3PO_lo_ treatment, the fold improvement in NP delivery to all TME subtypes was investigated (**Figure 7L-N**). The data reveal a lack of cancer cell-specific targeting upon 3PO_lo_ treatment. While the increase in relative levels of NP-bearing cancer cells was the highest for all 3 NP sizes, this was not a significantly better than the improvement for any of the other cell types. These data prove that 3PO treatment, and in particular 3POlo treatment is able to improve delivery of NPs to the tumor, and that this then results in a fairly even distribution of the relative increase per TME cellular subtype.

**Figure 7.**
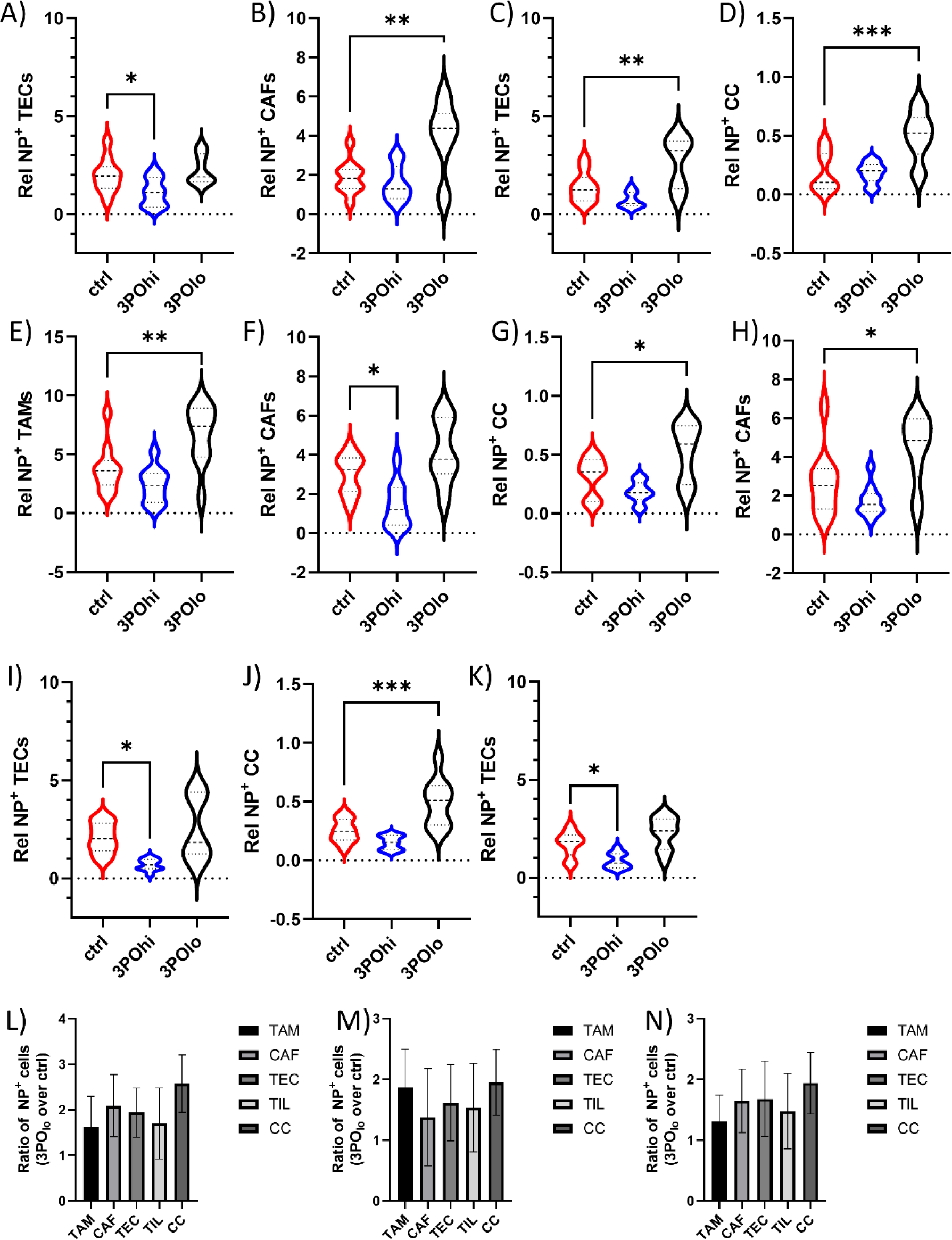
NP delivery to individual TME subtypes is increased upon 3PO_lo_ treatment. **a-e)** Violin plots showing the relative percentage of NP^+^ TME-related cell types expressed as the % of NP^+^ cells as determined by ImageStreamX Mark II expressed relative to the total gated cell type. Here, representative histograms are shown for those conditions where significant differences between control conditions and 3PO_lo_ or 3PO_hi_ are observed. Graph **a**) is for Au_10_, graphs **b-d**) are for Au_20_, graphs **e-g**) are for Au_40_, graphs **h-j**) are for Au_60_ and graph **k**) is for Au_80_. **l-n**) Histograms for **l)** Au_20_, **m)** Au_40_ and **n)** Au_60_ NPs which had displayed significant increases in NP delivery to tumor cells. The histograms represent the level of NP^+^ cells for every subtype expressed as the relative ratio of 3PO_lo_-treated animals versus control animals. Significant differences in all violin plots are indicated where appropriate (*n* = 15; *: p < 0.05; **: p < 0.01; ***: p < 0.001).

### 2.7 Evaluation of 3PO treatment on metastases

Several features of the TME have been described to foster malignancy and metastatic spread.(42–44) These include among others hypoxia, immune cell composition and vascularization. As 3PO_lo_ or 3PO_hi_ treatment affect both the level of hypoxia as well as the integrity of the tumor blood vessels, this raises the question whether the differences in NP extravasation could also suggest a possible increase in tumor cell intravasation? To analyze this, we performed a comparative study on the level of circulating tumor cells (CTC) in the blood, as well as the level of metastases observed in the lung, a common target for metastatic dissemination of Renca cells and renal cell carcinoma in general.(45) **Figure 8** reveals 3PO_lo_ and 3PO_hi_ had opposing effects on CTC counts in the blood, which translated into different effects on the level of tumor malignancy.

**Figure 8.**
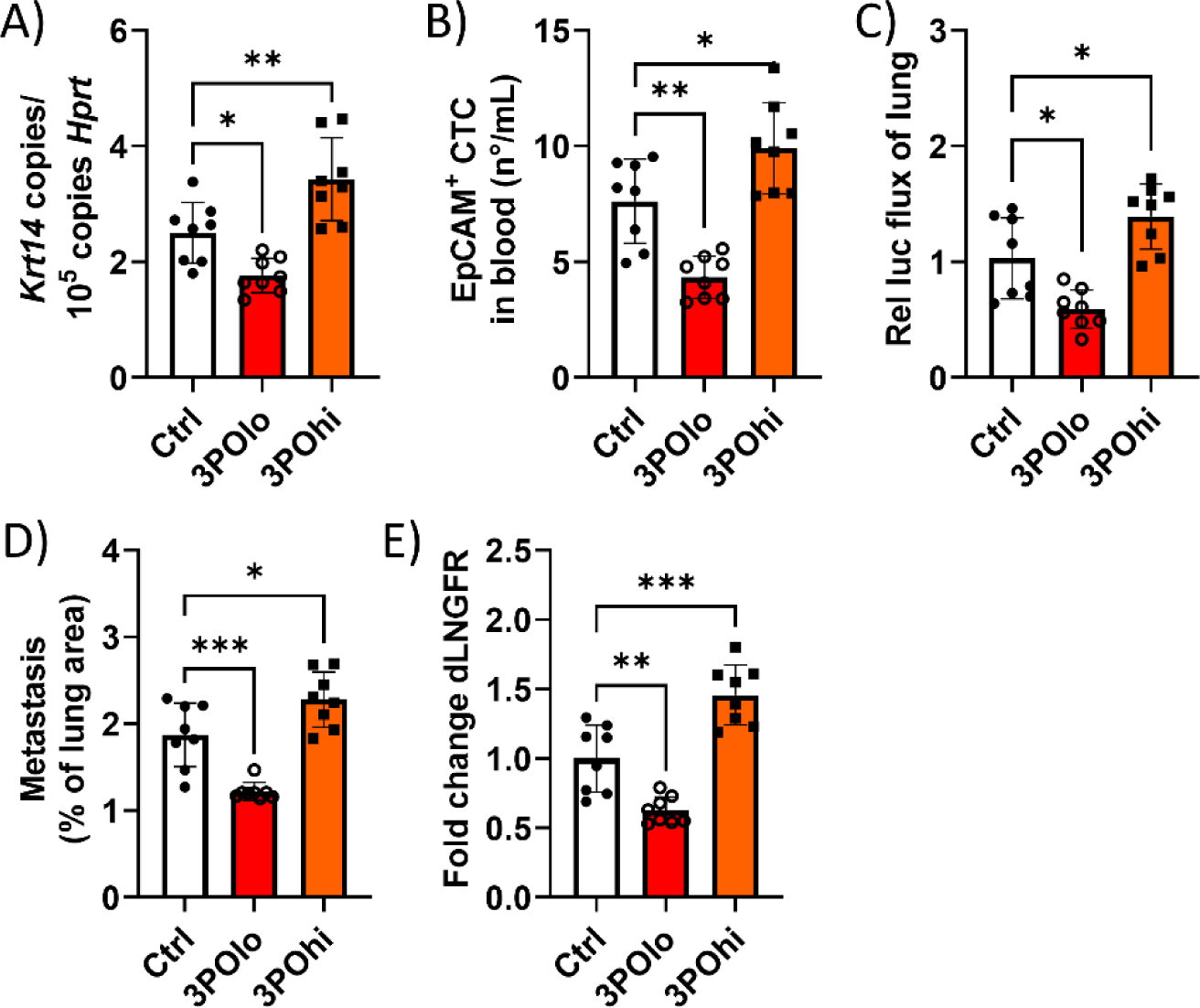
The effect of 3PO treatment on tumor metastasis. Histograms revealing the level of indicated markers obtained from lung tissue or blood samples derived from animals at 2 weeks following 3PO treatment or saline (control treatment). **a)** The level of *Krt14* expression relative to that of housekeeping gene *Hprt* determined by qPCR of blood samples taken at 14 days post 3PO treatment. **b)** The number of EpCAM^+^CD45^-^ cells detected in the blood at 14 days post 3PO treatment expressed as the number of EpCAM^+^ cells per mL of sample and determined by ImageStream analysis. **c)** Firefly Luciferase luminescence signal measured over the lung area in tumor-bearing mice before isolation of the tumors at 14 days post 3PO treatment. **d)** The area taken up by metastatic nodules in the lung tissue slice as determined by H&E staining relative to the area of the entire tissue slice. **e)** Expression of the truncated dLNGFR receptor that was introduced into the tumor cells and determined by qPCR of lung tissue and expressed relative to the level of untreated tumor-bearing mice. Significant differences between a treated group and untreated controls are indicated where relevant (p < 0.05: *, p < 0.01: **; p < 0.001: ***; p < 0.0001: ****) based on ANOVA testing using GraphPad 9 (*n* = 8).

The results obtained are in line with the findings above, where the increased permeabilisation of the tumor blood vessels upon 3PO_hi_ treatment result in increased tumor cell intravasation and final metastatic dissemination into the lungs. The normalized tumor blood vessels upon 3PO_lo_ treatment improved the barrier integrity and hereby impeded tumor malignancy. These results are also in line with literature data, where increased vessel permeabilisation due to 3PO or other means such as radiotherapy or hyperthermia can result in increased metastases.(46) Tumor vessel normalization by means of chemical of pharmacological means has been typically associated with a reduction in tumor malignancy.(47)

## 3 Discussion

The present work studied the effect of low and high concentrations of 3PO on tumor blood vessel maturation or permeabilisation and the implications of these treatments on delivery of NPs to solid tumors. Overall, the data reveal that, in line with literature results, 3PO_lo_ results in tumor vessel normalization, while 3PO_hi_ results in increased vessel permeabilisation.(16) In view of NP delivery to the tumor, 3PO_lo_ resulted in increased delivery efficacy, while 3PO_hi_ impeded NP delivery (**Scheme 1**).

**Scheme 1.**
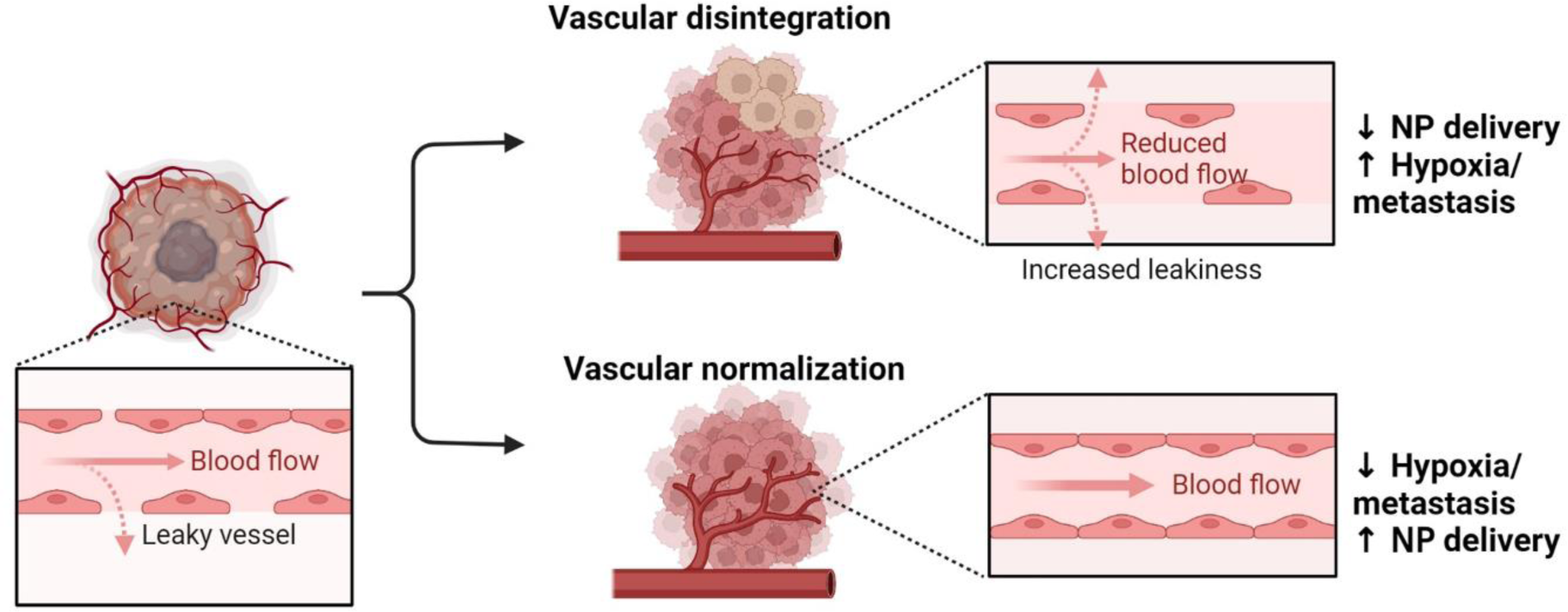
Schematic overview of the functional outcomes of vessel permeabilisation and normalization on NP delivery and tumor malignancy. Made by BioRender.

The latter is somewhat surprising, as both vessel normalization and permeabilisation have been associated with improved NP delivery efficacies.(11, 12, 23) The poor performance of 3PO_hi_ may be due to the way that 3PO results in vessel permeabilisation. High 3PO doses are given repeatedly over a long range of time, which was shown to significantly impact vessel integrity. However, at the same time, the loss in vessel integrity resulted in reduced perfusion of the tumor, which in turn will result in lower levels of NPs being perfused throughout the TME. The fact that perfusion was not halted altogether is likely reflected by the fact that only fast-growing, highly metabolically active TECs will be affected by 3PO_hi_, and more slow-growing vessels or vessels that were already present but are being redirected by the tumor itself as a result of vessel sprouting.(48) Therefore, any perfusion and most NPs as such, will therefore likely pass by the still intact, but more mature slow-growing vessels. As the total number of vessels in the tumor bed for 3PO_hi_-treated tumors is much lower than for control tumors, this likely explains the low NP delivery efficacy. Other studies that have observed higher NP delivery efficacies, may have used strategies (physical therapies such as radiotherapy or hyperthermia) that would be applied on the tumor shortly before or while the NPs are in circulation.(49, 50) Excessive vascular leakage has been shown to cause plasma escape and hemoconcentration. This was found to result in flow stasis and high interstitial pressure, which greatly hinder the extravasation of drugs and their movement to the tumor parenchyma.(11) This in itself supports our observation that extensive tumor vessel permeabilisation did not result in improved delivery, but rather reduced it. Another factor to take into account is the tumor model of choice. As levels of neoangiogenesis and TEC metabolism are quite variable between tumor types and even between different individuals bearing the same tumor, this can also play a major role on the impact of any vessel modification strategy on overall perfusion levels.

These data raise the question to what extent the use of blood vessel permeabilization approaches to improve NP delivery to solid tumors is warranted or necessary. To date, both normalization and permeabilization approaches are being explored for promoting NP extravasation in the tumor.(10, 12, 23) As extensive permeabilisation may impede NP delivery, it is imperative that the level of permeabilisation is well controlled and does not become excessive. However, the high heterogeneity of tumor vessel density and maturity in tumor models as well as in humans makes this very hard to achieve and this has been highlighted as one of the reasons why nanotherapeutics have underperformed.(32) In addition, the potential for increased malignancy associated with higher blood vessel permeability levels questions the safety of these approaches. While short-term attempts at vessel permeabilisation may be able to circumvent most of the tumor intravasation issues, physical methods such as radiotherapy have been linked to increased metastases levels in patients.(51) Great care must therefore be taken to design appropriate strategies for improved NP delivery.

Here, we suggest that tumor vessel normalization strategies would be a safer option and can at the same time result in improved NP delivery efficacies, although in this study, this was found to only be significant for NPs between 20 and 60 nm. For larger NPs, alternative means would be required in order to boost their delivery efficacy. In line with previous work, analyzing the tumor for the presence of so-called N-TECs, along with analysis of hypoxia, necrosis, and tumor vessel permeability and perfusion levels are good indicators to determine whether NPs can be used to target the tumor. Furthermore, as 3POlo was found to improve N-TEC levels, it would be interesting to see whether other strategies could promote N-TECs and hereby modulate NP delivery levels.

## 4 Materials and methods

### 4.1 Cell lines

Renca cell line was obtained from ATCC and cultured in RPMI 1640 supplemented with 10% fetal bovine serum (FBS), 1% penicillin/streptomycin, 1% non-essential amino acids and 1% sodium pyruvate (Gibco, Invitrogen, Merelbeke, Belgium). The cells were kept in a humidified 37°C incubator with a 5% CO_2_ environment. A lentiviral vector (LentiGuide) expressing a truncated form of low-affinity nerve growth factor receptor (dLNGFR) as a surface reporter protein was introduced by lentiviral transduction in Renca cells. Transduced cells were sorted to obtain a pure dLNGFR+ cell population and used for metastasis studies only. A lentiviral vector encoding a mutated (optimized for use in mouse models with no immunogenicity) click beetle luciferase (CBR2),(52) miRFP670 was introduced by lentiviral transduction in Renca cells. Transduced cells were sorted according to miRFP670 expression and miRFP670+ cells are hereby referred to as Renca-luc and used in metastasis studies.

### 4.2 Characterization of AuNPs

The core size of the AuNPs were characterized by transmission electron microscopy (TEM). Formvar film coated 400-mesh copper grids (Agar Scientific Ltd., England) were first glow-discharged to improve adsorption efficiency. Next, 10 µL of diluted NP sample (1/3 of each stock suspension) was dropped onto the grids and left to evaporate. The grids were examined using a JEM-1400 transmission electron microscope (JEOL, Japan) at accelerating voltage 80 keV. Hydrodynamic radii were measured on a PCS 100 spectrometer (Malvern, UK) at 25 °C, measuring the scattered light at a 90° angle. Samples were diluted with PBS until 0.5 mg/mL. The average value of 3 series of 10 different runs is given. Electrophoretic mobilities of the samples at identical dilutions were measured with a Zetasizer IIC instrument (Malvern, UK) at 25 °C. Nanoparticle tracking analysis was performed using a NanoSight NS300 system (Malvern, UK) using samples diluted in PBS containing 50% FBS at a final concentration of 5 µg/mL.

### 4.3 Tumor model

All animal experiments and research procedures were conducted in accordance with the declaration of Helsinki and EU Directive 2010/63/EU on the protection and welfare of animals used for scientific research. These experiments were approved by the Institutional Animal Care and Research Advisory Committee (KU Leuven) (ECD number: P203/2019) and were performed in accordance with the institutional and national guidelines and regulations. Female Balb/c mice, 5 weeks old with body weights of 18-25g, were purchased from Charles River (Wilmington, MA, US) and housed in a specific pathogen-free environment.

1 x10^6^ Renca Firely-Luciferase and GFP-positive (Renca Luc/GFP) cells were injected subcutaneously in the right flanks of the mice to asses subcutaneous tumor growth. Tumor volumes were measured with calipers and calculated using the formula V=∏ x ((d^2^xD)/6), where d is the minor tumor axis and D the major tumor axis.

### 4.4 Pharmacological treatments

When the tumor sizes reached to 80 mm^3^ animals were treated with a PFKFB3 inhibitor 3-(3-pyridinyl)-1-(4-pyridinyl)-2-propen-1-one) (3PO) (Sigma-Aldrich, Merck, Overijse, Belgium). The animals received intraperitoneal (i.p.) injections 3x a week (consecutive days) during 2 weeks of 25 mg/kg 3PO or 75 mg/kg 3PO or dimethyl sulfoxide (DMSO) (Sigma-Aldrich, Merck, Overijse, Belgium) as control. The treatment time-points were as following with the start of the treatment marked as day 1: day1, day2, day3, day 8, day9, and day10 (see Figure 2).

### 4.5 Tumor vascular perfusion and permeability

48h after the last pharmacological treatment of the animals, the tumor vascular perfusion and permeability was determined by injecting intravenous (250 ug/ml) bovine serum albumin functionalized with Alexa Fluor 555 (BSA-AF555, Thermo Fisher Scientific, Ghent, Belgium) and measured *in vivo* using the IVIS Spectrum (Perkin Elmer, Life Sciences, Zaventem, Belgium) during 90 minutes with sequential measurements every 3 minutes (10 sec, medium binning, excitation 500 nm 4nd emission 580 nm). Measurements occurred immediately upon intravenous administration of BSA.

### 4.6 Animal experiment

Mice were injected intravenously with AuNPs (80 µg Au/mouse in 100 µL) 24h following the perfusion/permeability assay, with sizes ranging from 10, 20, 40, 60 or 80 nm. The animals were sacrificed with Dolethal (Pentobarbital Sodico) (200 mg/ml, Vetoquinol, Aartselaar, Belgium) 72h after the AuNPs injections. The tumor and all the major organs (heart, lungs, spleen, kidney and liver) were fixed in 4% paraformaldehyde (PFA, Klinipath, VWR, PA, USA).

### 4.7 *In vivo* hypoxia detection

Animals were injected intravenously 24h after the last pharmacological treatment with Hypoxisense 680 (2 nmol/100 uL per mouse, Perkin Elmer, Life Sciences, Zaventem, Belgium), and the Hypoxisense 680 was measured *in vivo* 24h post-injection using the IVIS Spectrum (Perkin Elmer, Life Sciences, Zaventem, Belgium). The animals were sacrificed 72h after the Hypoxisense 680 measurement with Dolethal (Pentobarbital Sodico) (200 mg/ml, Vetoquinol, Aartselaar, Belgium) and the tumors were fixed in 4% paraformaldehyde (PFA, Klinipath, VWR, PA, USA). Only histological procedures were performed on these tumors.

### 4.8 Histology, immunostainings and morphometric analysis

Mouse tissues were dehydrated after being fixed for minimum 48h in 4% PFA at 4°C. The tissues were after subsequently embedded in OCT compound (Sakura-Finetek, CA, USA) and frozen at −80°C. Tissue slices of 10 µm were cut using the Cryostar NX70 (Thermo Fisher Scientific, Ghent, Belgium) and placed on glass microscope slides (VWR, PA, USA). For morphometric analyses, optical fields (40x magnification) of the whole sections were taken by the high content screening microscope Nikon-Marzhauser Slide Express (Märzhäuser Wetzlar GmbH & Co. KG, Wetzlar, Germany) or Vectra Polaris multispectral imaging system (Perkin Elmer, Life Sciences, Zaventem, Belgium) and analysed using QuPath. The images were automatically stitched together to cover the entire slide and were saved as .OME TIFF of .qptiff file format for further analysis.

#### 4.8.1 Hematoxylin and eosin staining for determination of tumor necrosis and tissue damage

Hematoxylin and eosin (H&E) staining was performed on 10 µm thick OCT-embedded frozen tissues obtained as described above. The tumor slides where first air-dried without dehydration for 30’ at RT and washed with 1X Phosphate buffered saline (PBS, Gibco, Thermo Fisher Scientific, Ghent, Belgium) for 5’. The slides were then stained protected from light with hematoxylin (Sigma-Aldrich, Merck, Overijse, Belgium) for 3’, washed with deionized water obtained from a Milli-Q system (MQ; Millipore, France) water for 5’, incubated for 1’ in ethanol 80% (0.15% HCl), washed for 1’ with MQ and incubated for 30’’ with ammonium-containing water, followed by washing for 5’ with MQ water and 95% ethanol for 1’. The tumor slides were incubated protected from light for 1’ in eosin (Sigma-Aldrich, Merck, Overijse, Belgium) and dehydrated after in 95% ethanol for 5’, twice in 100% ethanol for 5’ each followed by washing in Xylene (Sigma-Aldrich, Merck, Overijse, Belgium) twice for 5’ each. Finally, the samples were mounted with DPX mounting medium (Merck, Overijse, Belgium).

Tumor necrosis was then expressed as the percentage of the total tumor area as determined on H&E-stained sections. For this, the images were loaded as Brightfield (H&E) images, and a thresholder was created to classify pixels for eosin (tumor tissue) or background signal. This was followed by creating a thresholder for classifying pixels for hematoxylin signal, to discriminate healthy from necrotic tissue. The tumor area was then calculated based on the eosin stain, while the area of the healthy tissue is then calculated based on the hematoxylin stain. The relative level of necrosis is then calculated as follows: ((eosin area – hematoxylin area)/eosin area)*100 and expressed as percentage for all tissue sections and all tumors. Detailed procedures of the analysis method are described in Izci et al.(13)

#### 4.8.2 Picrosirius red staining for determination of tumor extracellular matrix level

The tumor slides were air-dried without dehydration for 30’ at RT and washed with 1X PBS (Gibco, Thermo Fisher Scientific, Ghent, Belgium) for 5’. The tumor samples were stained with Picrosirius red (PSR, Abcam, Cambridge, UK) and Fastgreen (0.1g FCF in acidified PSR solution, Sigma-Aldrich, Merck, Overijse, Belgium) for 1h at 4°C and rinsed twice with 0.5% acidified water followed by washing in Ethanol absolute for 30” one time and for 5’ two times. The samples were cleaned twice in Xylene for 5’ and mounted with DPX mounting medium (Merck, Overijse, Belgium).

Tumor extracellular matrix content was expressed as the percentage of the total tumor area. For analysis in QuPath, the images were loaded as Brightfield (H-DAB) images, and the images were preprocessed and stain vectors were estimated and adjusted to represent the PSR and Fastgreen signal, respectively (see Izci et al. for detailed procedure(13)). Using the combined colours, a threshold was created to classify pixels for tumor tissue or background signal. This was followed by creating a threshold for classifying pixels for PSR signal, to discriminate collagen fibres from other tissue. The tumor area was then calculated based on the combined stain, while the area of the collagen is then calculated based on the PSR stain. The relative level of necrosis is then calculated as follows: (PSR area/total area)*100 and expressed as percentage for all tissue sections and all tumors.

#### 4.8.3 Immunohistochemistry

The tumor samples were air-dried at RT for 30’ without dehydration and were washed in 1X PBS for 5’. The samples were fixed in 100% cold MeOH for 6’ at −20°C, washed for 5’ in 1XPBS, and incubated for 15’ with Proteinase K (1:500, in 1XPBS, Promega B.V., Leiden, The Netherlands) at 37°C. After washing the slides 5’ with 1XPBS, the slides were blocked for 1h at RT with 1XPBS + 10% normal goat serum (NGS, 60.0 mg/mL, Thermo Fisher Scientific, Ghent, Belgium) and 1%FBS and washed again twice with 1XPBS for 5’ each. The samples were blocked with Avidin (0.001%, Avidin from egg white, Sigma-Aldrich, Merck Chemicals, Overijse, Belgium) for 20’, followed by washing with 1XPBS two times for 2’ each and blocked again with Biotin (0.001%, Sigma-Aldrich, Merck, Overijse, Belgium) for 20’ and washed twice for 2’ in 1XPBS each. The samples were incubated with anti-CD31 antibody (1:25, in 1XPBS + 1%NGS, Abcam, Cambidge, UK) overnight at 4°C.

After leaving the samples for 20’ at RT, the samples were washed twice in 1XPBS for 5’ each, blocked for 20’ with hydrogen peroxidase (3%, Alexa Fluor 594 Tyramide SuperBoost Kit, Invitrogen, Thermo Fisher Scientific, Ghent, Belgium), washed twice in 1XPBS for 5’ each and incubated with Goat anti-Rat-Biotin (1:300, in 1XPBS + 1%NGS, Jackson ImmunoResearch Europe Ltd, Ely, UK) at RT for 1h.

The samples were washed twice in 1XPBS for 5’ each, incubated with streptavidin-HRP (1:150, in 1XPBS + 1%NGS, Invitrogen, Thermo Fisher Scientific, Ghent, Belgium) for 30’ at RT, washed again two times in 1XPBS 5’ each and incubated with Alexa Fluor Tyramide 594 (1:100, Alexa Fluor 594 Tyramide SuperBoost Kit, Invitrogen, Thermo Fisher Scientific, Ghent, Belgium) + hydrogen peroxidase 3% (1:100, Alexa Fluor 594 Tyramide SuperBoost Kit, Invitrogen, Thermo Fisher Scientific, Ghent, Belgium) + Tris Buffer HCl pH 7,4 (1:1) for 10’ at RT. The reaction was stopped using Stop Reagent (1:11, in 1XPBS, Alexa Fluor 594 Tyramide SuperBoost Kit, Invitrogen, Thermo Fisher Scientific, Ghent, Belgium) for 2’ at RT, washed three times in 1XPBS for 5’ and incubated with anti-neural/glial antigen 2 (NG-2, 1:100, in 1XPBS + 1%NGS, Abcam, Cambidge, UK), overnight at 4°C.

##### Anti-CD31 and anti-NG2 co-staining

The samples were washed twice in 1XPBS for 5’ each, incubated with Goat Anti-Rabbit IgG secondary antibody poly HRP (1:1, Alexa Fluor 488 Tyramide SuperBoost Kit, goat anti-rabbit IgG, Invitrogen, Thermo Fisher Scientific, Ghent, Belgium) 1h at RT, washed twice in 1XPBS for 5’ each, incubated with Alexa Fluor Tyramide 488 (1:100, Alexa Fluor 488 Tyramide SuperBoost Kit, goat anti-rabbit IgG, Invitrogen, Thermo Fisher Scientific, Ghent, Belgium) + hydrogen peroxidase 3% (1:100, Alexa Fluor 488 Tyramide SuperBoost Kit, goat anti-rabbit IgG, Invitrogen, Thermo Fisher Scientific, Ghent, Belgium) + Tris Buffer HCl pH 7,4 (1:1) for 10’ at RT, incubated after for 2’ at RT with Stop Reagent (1:11, in 1XPBS, Alexa Fluor 488 Tyramide SuperBoost Kit, goat anti-rabbit IgG, Invitrogen, Thermo Fisher Scientific, Ghent, Belgium), washed three times in 1XPBS for 5’ each and incubated for 10’ at RT with Hoechst (1:1000, in 1XPBS, Invitrogen, Thermo Fisher Scientific, Ghent, Belgium). Finally, the samples were washed twice in 1XPBS for 5’ each, mounted with Fluoromont (Sigma-Aldrich, Merck Chemicals, Overijse, Belgium), air-dried for 30’ with dehydration and sealed the cover slides with transparent nail polish.

<u>Tumor vessel area</u> was analysed by immunostaining CD31, which is a marker for endothelial cells, where the total CD31^+^ area was expressed as the percentage of the total tumor area. For analysis in QuPath, the entire image was annotated as a field (tumor tissue), where the number of cells were calculated using “cell detection” based on DAPI signal. Positive cell detection was then performed by setting threshold for the AF594 signal. The total number of endothelial cells then were determined as well as the total area of CD31^+^ positive cells compared to other cells. The relative level of CD31^+^ vessel density is then calculated as follows: ((CD31 area)/total area)*100 and expressed as percentage for all tissue sections and all tumors (please see Izci et al for detailed analysis protocols(13)).

<u>Tumor vessel maturation</u> was assessed by the immunostaining for NG-2, which is a pericyte marker together with immunostaining for CD31. For analysis in QuPath, the entire image was annotated as a field (tumor tissue), where the number of cells were calculated using “cell detection” based on DAPI signal. Positive cell detection was then performed by setting threshold for the AF594 signal (CD31 positive) or AF488 signal (NG2 positive). For analysis, the CD31^+^ cells and NG2^+^ cells were artificially dilated 2-fold and any enlarged cell (typically the size of at least 2 cells), comprising both CD31 and NG2 signal is determined as a mature vessel. The total number of mature endothelial cells then were determined as well as the total area of NG2^+^CD31^+^ double positive cells compared to total number of CD31^+^ endothelial cells. The relative level of NG2^+^CD31^+^ vessel density is then calculated as follows: ((NG2^+^CD31^+^ area)/CD31^+^ area)*100 and expressed as percentage for all tissue sections and all tumors.

### 4.9 RNAscope analysis

RNA in situ hybridization for mouse *Plvap* (440221-C1), mouse *CD276* (590091-C3) and mouse *CD31* (471481-C2) was performed according to the manufacturer’s instructions (Advanced Cell Diagnostics). Briefly, a total of 10 sections were selected, 5 sections tissues obtained from normal tumor samples (control mice) and 5 sections of tumors obtained from tumor-bearing mice treated with 3PO_lo_. Of the tissue samples, 10 μm paraformaldehyde-fixed, OCT-embedded frozen intestinal tumour sections were pretreated with heat in the retrieval reagent and protease III before hybridization with the target oligonucleotide probes. Preamplifier, amplifier and alkaline-phosphatase-labelled oligonucleotides were then hybridized sequentially. HRP signal was developed using Opal520 (Akoya Biosciences, FP1487001KT) for the CD31 probe, Opal570 (Akoya Biosciences, FP1488001KT) for the *Plvap* probe and Opal620 (Akoya Biosciences, FP1495001KT) for the *CD276* probe. Quality control was performed to assess RNA integrity with probes specific to ubiquitously expressed household genes PolR2A RNA (320881-C1), PPIB RNA (320881-C2), UBC RNA (320881-C3) and for background staining with a probe specific to bacterial dapB RNA (320871). Samples were counterstained with DAPI nuclear counterstain and imaged using the Vectra Polaris multispectral imaging system (Perkin Elmer, Life Sciences, Zaventem, Belgium) and analysed using QuPath. Specific fluorescent signal for *CD31*, *Plvap* and *CD276* was identified as green, red and far-red punctate dots, respectively. For analysis, *CD31*^+^ cells were identified as cells with green positive dots and in these cells, the presence of *CD276* and *Plvap* (single or both) in these cells was determined and expressed relative to the total amount of *CD31*^+^ cells.

### 4.10 Single cell analysis of NP uptake by image-based cytometry

The tumor samples were dissociated into single cells using GentleMACS tissue dissociator and its kit (Miltenyi Biotec, Gladbach, Germany). The tumor samples were cut into small pieces, transferred in gentleMACS C-tubes containing RPMI/DMEM media and kit enzymes (tumor dissociation kit, Miltenyi Biotec, Gladbach, Germany) and were broken down using the gentleMACS. After the tumors were processed, the gentleMACS C-tubes were centrifuged 30’ on 1,5 rpm, the content of the gentleMACS C-tube was passed first through 70 um, followed by 40 um strainer and was centrifuged for 7’ on 300g. The pellet was lysed using RBC lysis buffer for exactly 2’, centrifuged for 5’ on 300g and resuspended in 1 mL media, which was after added slowly on top of 1mL FBS to form a layer of media on top of the FBS. After centrifugation for 5’ on 100g, the single tumor cells inside the media sank to the bottom of the FBS and the tumor cells were separated from the debris. The supernatant was removed, the cells were washed with 1XPBS, centrifuged for 5’ on 1.4 rpm and were incubated with Fc Blocker (1:100, in 1XPBS + 1%FBS, Thermo Fisher Scientific, Ghent, Belgium) for 30’ on ice. The cells were washed with 1XPBS+ 1%FBS, centrifuged for 5’ on 1.4 rpm and incubated with two different antibody cocktails for 1h on ice, protected from light, where all the antibodies were diluted in 1XPS+ 1%FBS. The following antibodies were used in the first cocktail; anti-CD45 FITC (1:100, Thermo Fisher Scientific, Ghent, Belgium), anti-90.2 APC (2:100, Thermo Fisher Scientific, Ghent, Belgium), anti-CD3 AF610 (2:100, Thermo Fisher Scientific, Ghent, Belgium) and anti-CD144 PE (2:100, Thermo Fisher Scientific, Ghent, Belgium), and the following two antibodies in the second cocktail; anti-CD24 APC (3:100, Thermo Fisher Scientific, Ghent, Belgium) and anti-F4/80 FITC (3:100, Bio-Rad Laboratories, Temse, Belgium). The single cells were washed with 1XPBS+1%FBS, resuspended in 1XPBS and transported in eppendorfs to the image-based cytometer Imagestream Mark II Imaging flow cytometer (Merck, Overijse, Belgium). Measurements were done by acquiring approximately 1×10^5^ single cells per sample. The images were acquired using the 40x objective with the darkfield (780 nm laser) at 0.3 mW in order to reduce scatter light, while enabling darkfield-based detection of AuNPs inside cells. For the first cocktail, laser intensities were set at 1.00mW (488 nm), 20.00 mW (561 nm) and 50.00 mW (642 nm) and for the second cocktail, these were set at 1.00 mW (488 nm) and 20 mW (642 nm).

For analysis, iDEAS software (Amnis Corporation, USA) was used, followed by FCS Express 7.0 for visualization. First, focused and single cell were selected and gated, after which cell selections were gated based on the different markers used: tumor cells were defined as CD45^-^CD24^+^, tumor-associated macrophages as F4/80^+^, leukocytes as CD45^+^, endothelial cells as CD144^+^ and cancer associated fibroblasts as CD45^-^CD90.2^+^. For every cell type, darkfield images were taken and signal obtained in the darkfield channel were analysed for the different cell types, using bright intensity projection. Please see Izci et al. for detailed gating and analysis protocols.(13)

### 4.11 Inductively coupled mass spectrometry

#### 4.11.1 Instrumentation

(Ultra-)trace element determination of Au was carried out using an Agilent 8800 ICP-MS/MS instrument (ICP-QQQ, Agilent Technologies, Japan). The sample introduction system comprises a concentric nebulizer (400 µL min^-1^) mounted onto a Peltier-cooled (2 °C) Scott-type spray chamber. This instrument is equipped with a tandem mass spectrometry configuration consisting of two quadrupole units (Q1 and Q2) and a collision/reaction cell (CRC) located in-between both quadrupole mass filters (Q1-CRC-Q2). All measurements were performed in MS/MS mode (on-mass approach) with the collision/reaction cell (CRC) operated in “vented” (no gas) mode.

#### 4.11.2 Reagents and standards

For ICP-MS/MS analysis, only high-purity reagents were used. Ultra-pure water (resistivity 18.2 MΩ cm) was obtained from a Milli-Q Element water purification system (Millipore, France). Pro-analysis purity level 14 M HNO_3_ (Chem-Lab, Belgium) further purified by sub-boiling distillation and ultra-pure 9.8 M H_2_O_2_ (Sigma Aldrich, Belgium) were used for sample digestion. Appropriate dilutions of 1 g L^-1^ single element standard solutions of Au and Tl (Inorganic Ventures, USA) were used for method development, optimization, and calibration purposes. For quantitative element determination of Au, external calibration was relied on as calibration approach (0, 0.1, 0.25, 0.5, 1.0 and 2.5 µg L^-1^ Au), with Tl (2 µg L^-1^) as internal standard.

#### 4.11.3 Samples and sample preparation

The samples were digested *via* acid digestion in Teflon Savillex® beakers, which had been pre-cleaned with HNO_3_ and HCl and subsequently rinsed with Milli-Q water. A mixture of 1 mL of 14 M HNO_3_ and 0.5 mL of 9.8 M H_2_O_2_ was added to each sample (mass ranging between 1 and 770 mg) and the procedure was completed after heating at 110 °C on a hot plate for approximately 18 h. Prior to ICP-MS/MS analysis, the digests were appropriately diluted (between 10- and 2000-fold dilution) with Milli-Q water. To avoid contamination, only metal-free tubes were used for standard and sample preparation (15 mL polypropylene centrifuge tubes, VWR, Belgium). Tl was added to all samples and standards to correct for instrument instability, signal drift and matrix effects.

### 4.12 NP biodistribution analysis per parameter

The effect of the different parameters measured (NP diameter, tumor size, ECM density, necrotic area, vessel perfusion, vessel permeability, blood vessel area, blood vessel maturity, blood vessel-ECM, number of TAMs, number of CAFs, number of endothelial cells, number of lymphocytes) on NP distribution (efficacy of NP accumulation in total tumor area and efficacy of NP accumulation in tumor cells) is calculated using Uniform Manifold Approximation and Projection (UMAP) based dimensionality reductions (FlowExpress 7.0) to assess which parameters caused the biggest influence on NP tumor accumulation.

### 4.13 Metastases analysis

Renca-dLNGFR, or Renca-luc were used and implanted subcutaneously, after which they got treated with 3PO_lo_, 3PO_hi_ or saline as described in section 4.3. At 14 days following 3PO treatment, lung tissue was collected after perfusion of animals with a saline through the right ventricle and metastasis were assessed by hematoxylin-eosin staining on frozen tissue sections. For *ex vivo* quantification of cancer cell burden, at 14 days following 3PO treatment, lung tissue was collected after perfusion of animals with a saline through the right ventricle and either bioluminescence was measured in an IVIS Spectrum, snap-frozen (to assess dLNGFR expression by qRT-PCR), or fixed in 2% PFA and frozen in OCT, metastasis were assessed by hematoxylin-eosin staining on frozen tissue sections.

For analysis of circulating tumor cells (CTCs), blood samples were collected and centrifuged in heparin-containing tubes to separate plasma from serum (15 min at 3500 rpm). The serum was then depleted from CD45^+^ cells using magnetic separation beads (StemCell Technologies), and the remaining samples were split into 2 fractions, where one was evaluated for expression of epithelial marker *Krt14* (not present in blood cells), and household gene *Hprt* by qPCR. The other fraction was used to stain with anti-CD45 (FITC, Thermo Fisher) and anti-EpCAM (PE-Texas Red, Thermo Fisher) antibodies and analyzed using ImageStream analysis, where CTCs were defined as EpCAM^+^CD45^-^ cells.

### 4.14 Blood biochemistry

Upon isolation of the tumors and main organs for analysis, blood samples were also collected of control mice bearing Renca tumors but without any AuNPs or mice Balb/c mice with Renca tumors having received a bolus of AuNPs. Blood samples were collected retroorbitally following animal sacrifice (200 µl/animal), and samples were collected and centrifuged in heparin-containing tubes to separate plasma from serum (15 min at 3500 rpm). Next, 75 µl serum was added on analysis discs (Samsung Comprehensive test 16V) enabling analysis of 16 different markers using the Samsung PT10V chemistry analyzer (SCIL Animal care company GmbH, Viernheim, Germany). The following markers were analyzed: glucose, urea, creatinine, urea/creatinine ratio, phosphates, calcium, total protein, albumin, globulin, albumin-globulin ratio, alanine aminotransferase, alkaline phosphatases, bilirubin, cholesterol, triglycerides and amylase.

### 4.15 Statistical analysis

All statistical analyses were performed using GraphPad 9.0 statistical analysis software. To determine significant differences between groups, 2-way ANOVA tests were performed with Tukey post-hoc test, unless otherwise indicated in the corresponding text. The levels of significance and number of independent repeats are indicated with every data point given.

## 5 Declarations

### 5.1 Ethics approval and consent to participate

All animal experiments and research procedures were conducted in accordance with the declaration of Helsinki and EU Directive 2010/63/EU on the protection and welfare of animals used for scientific research. These experiments were approved by the Institutional Animal Care and Research Advisory Committee (KU Leuven) (ECD number: P203/2019) and were performed in accordance with the institutional and national guidelines and regulations.

### 5.2 Consent for publication

All authors have approved this manuscript for publication.

### 5.3 Availability of data and materials

All data needed to evaluate the conclusions in the paper are present in the paper and/or the Supplementary Materials. The raw data on tumor characterization and NM delivery efficacy can be provided by the corresponding author pending scientific review and a completed material transfer agreement. Requests for the raw numerical data should be submitted to.

### 5.6 Authors’ contributions

All authors contributed to the design, data retrieval, and writing of the manuscript.

## 5.7 Acknowledgements

Not applicable

## Supporting information

**Supplementary Figure S1.**
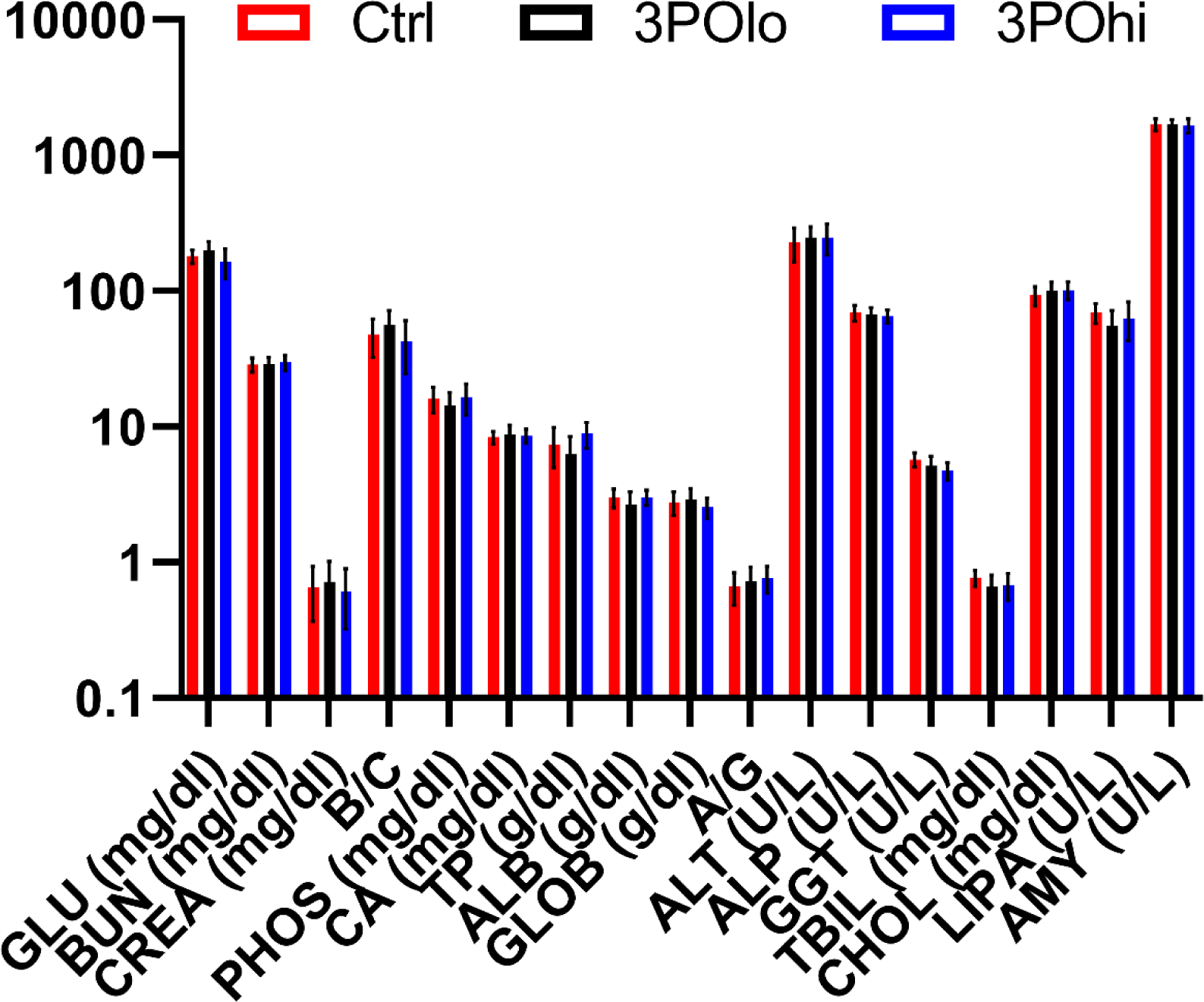
Blood biochemistry results indicate no toxicity of 3PO treatments. Histograms showing blood biochemistry results of Renca-bearing control mice (red bars) or Renca-bearing mice exposed to 3PO_lo_ (black bars) or 3PO_hi_ (blue bars) All data are expressed as mean ± SEM (*n* = 6). The following markers are studied: glucose (GLU), blood urea nitrogen (BUN), creatinine (CREA), ratio of urea over creatinine (B/C), phosphates (PHOS), calcium (CA), total protein (TP), albumin (ALB), globuline (GLOB), ratio of albumin over globuline (A/G), alanine aminotransferase (ALT), alkaline phosphatase (ALP), gamma-glutamyl transpeptidase (GGT), total bilirubin (TBIL), cholesterol (CHOL), triglycerides (LIPA), alpha-amylase (AMY).

**Supplementary Figure S2.**
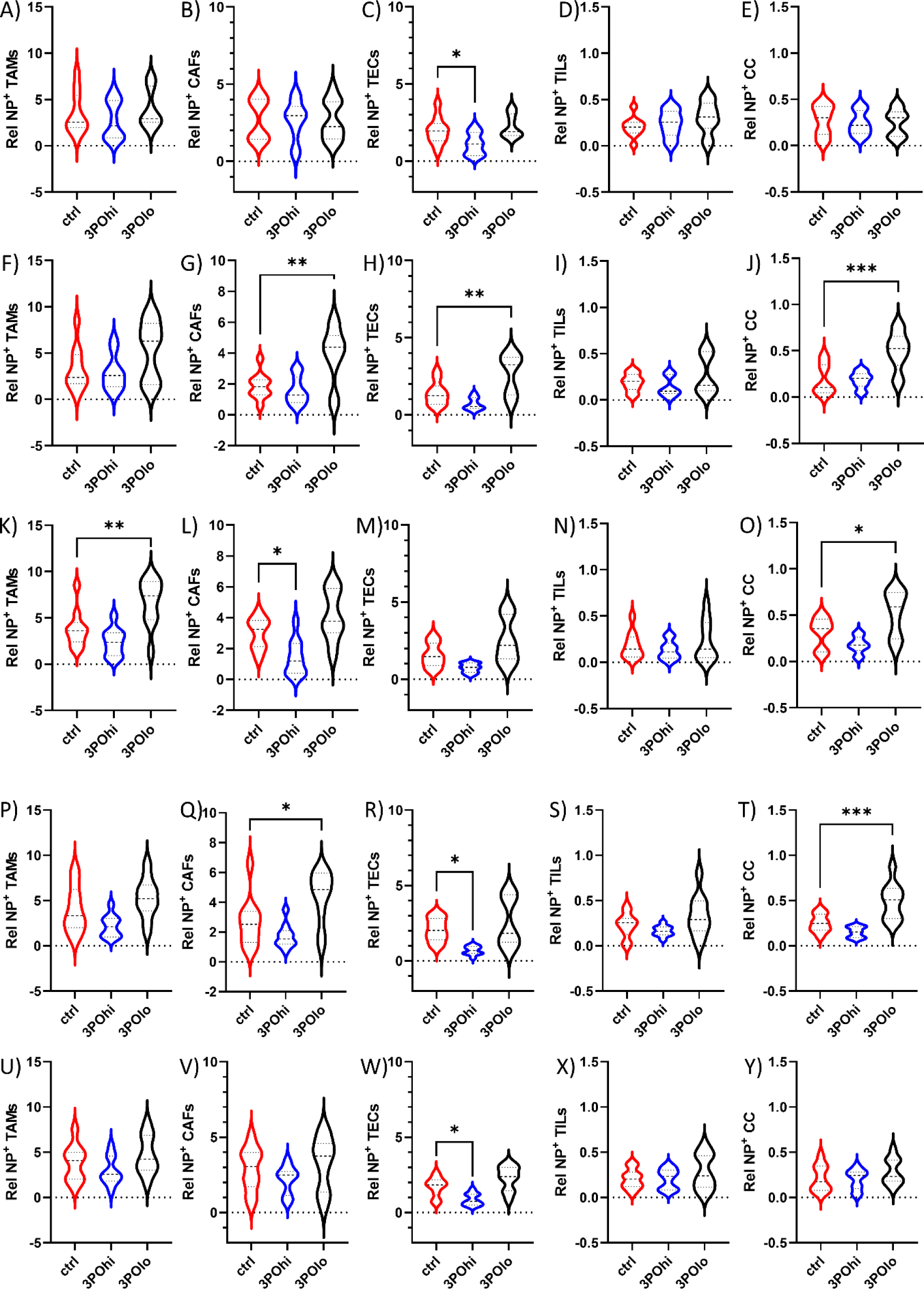
The effects of 3POlo and 3POhi treatment on NP delivery to TME-specific cell types. Violin plots showing the relative percentage of NP^+^ TME-related cell types expressed as the % of NP^+^ cells as determined by ImageStreamX Mark II expressed relative to the total gated cell type. Here, representative histograms are shown for those conditions where significant differences between control conditions and 3PO_lo_ or 3PO_hi_ are observed. Graphs **a-e)** are for Au_10_, graphs **f-j)** are for Au_20_, graphs **k-o)** are for Au_40_, graphs **p-t)** are for Au_60_ and graphs **u-y)** are for Au_80_. Significant differences in all violin plots between a 3PO-treated group and the controls are indicated where appropriate (*n* = 5; *: p < 0.05; **: p < 0.01; ***: p < 0.001).

